# Advancing Luciferase Activity and Stability beyond Directed Evolution and Rational Design through Expert Guided Deep Learning

**DOI:** 10.1101/2025.08.25.672183

**Authors:** Spencer Gardiner, Joseph Talley, Christopher Haynie, Joshua Ebbert, Corbyn Kubalek, Matthew Argyle, Deon Allen, William Heaps, Tyler Green, Dallin Chipman, Bradley C Bundy, Dennis Della Corte

## Abstract

Engineered luciferases have transformed biological imaging and sensing, yet optimizing NanoLuc luciferase (NLuc) remains challenging due to the inherent stability-activity trade-off and its limited sequence homology with characterized proteins. We report a hybrid approach that synergistically integrates computational deep learning with structure-guided rational design to develop enhanced NLuc variants that improve thermostability and thereby activity at elevated temperatures. By systematically analyzing libraries of engineered variants, we established that modifications to termini and loops distal from the catalytic center, combined with preservation of allosterically coupled networks, effectively enhance thermal resilience while maintaining enzymatic function. Our optimized variants – notably B.07 and B.09 – exhibit substantial thermostability enhancements (increases of 4.2 °C and 5.2 °C at 50 % solubility), leading to increased activity at elevated temperatures (320 % and 370 % of wild-type at 55 °C). These variants maintain NLuc’s pH tolerance and retain improved activity with the alternative substrate coelenterazine. Molecular dynamics simulations and protein folding studies elucidate how these mutations favorably modulate conformational landscapes without perturbing substrate binding architecture. Beyond providing superior tools for bioluminescence applications, our integrated methodology establishes a broadly applicable framework for engineering enzymes where traditional homology-based approaches fail, and stability-activity constraints present formidable barriers to improvement.

## Introduction

Bioluminescence, the production and emission of cold light by living organisms, provides a distinct optical readout for biological processes, distinguishing it from fluorescence, combustion, or electrical illumination [1, 2]. Luciferases, enzymes responsible for this effect, have become indispensable tools in protein–protein and protein–ligand interaction studies [3–6], gene regulation [6], protein stability monitoring [7], and bioimaging [8, 9]. Among these, bioluminescence resonance energy transfer (BRET)-based sensors have expanded the capabilities of real-time, noninvasive imaging, offering high signal-to-background ratios and detection sensitivity in deep tissues [10, 11].

Despite their utility, natural luciferases often suffer from poor folding, low activity, and susceptibility to proteolytic degradation in mammalian cells, limiting their application [12–14]. Engineering efforts have sought to overcome these challenges by *de novo* design or optimizing existing luciferases for increased brightness, stability, and compatibility with synthetic luciferins that offer desirable photophysical properties and higher substrate specificity [12]. *De novo* design has successfully created several luciferases that have high thermal stability (T_melt_ > 95 °C); however, these enzymes are substantially less luminescent than naturally derived luciferases [12, 15].

While Firefly and Renilla luciferases have been the traditional standards in bioassays and molecular imaging due to their well-characterized properties, the discovery and optimization of a deep-sea shrimp luciferase yielded the NanoLuc (NLuc) enzyme, a compact (19.1 kDa), stable luciferase with over 150-fold increase in luminescence relative to its natural precursors [7, 16].

Its monomeric nature and unbiased distribution in cells make it a versatile platform for numerous applications, including bioluminescent imaging and biosensing [7]. Its split variants (NanoBiT) have aided protein interaction studies by providing a dynamic and reliable reporter system [10, 17]. However, NLuc exhibits low thermal stability, limiting its use in high-temperature applications, such as wastewater monitoring [18], food processing [19], and fermentation control [20].

Our recent work showed that deep learning–based sequence optimization using BayesDesign can generate NLuc variants with markedly enhanced thermostability [21]. However, these variants suffered substantial loss of bioluminescent activity, underscoring a key limitation: stability-focused computational design alone is inadequate for optimizing functional performance in bioluminescent enzymes. Established rational design and deep learning methodologies frequently leverage homology guided insights to preserve activity [22–26]. However, NLuc shares limited sequence homology with known proteins [27], rendering homology-based design approaches, such as those employed by Sumida et. al. [28] ineffective for this enzyme. Additionally, due to the approximately 10^220^ possible mutations for NLuc, experts on the structure and function of NLuc should provide guardrails to limit which residues can be mutated to preserve key sites.

Structural studies have provided critical insights into NLuc structure and function, revealing a homotropic negative allosteric mechanism that complicates mutational improvements – changes distant from the active site can inadvertently disrupt activity. Rational design approaches based on these insights have successfully enhanced NLuc activity, however stability of most active mutants was not previously reported [29].

Here, we demonstrate that integrating rational design with deep learning can overcome the stability-activity trade-off and develop superior NLuc variants with enhanced stability, leading to increased activity at elevated temperatures. In this hybrid approach, structural and mechanistic insights are used to identify potentially mutable regions of the protein, which are then mutated using deep learning tools. Here, we report how this initial application enabled discovery of novel NLuc variants with enhanced properties that could significantly improve their utility in bioimaging, biosensing, and other bioluminescence-based applications.

## Results & Discussion

### Screening campaign

In this work, we demonstrate the capabilities of rational design-guided deep learning by overcoming the stability-activity trade-off historically observed with the NLuc protein. Recent efforts to optimize NLuc activity [29] demonstrated that rational design can go beyond directed evolution, although we found melting temperature to decrease by 8.6 °C (see Figure 6, variant A.01). Conversely, our previous work [21] employed BayesDesign, a deep learning approach, to engineer highly stable NLuc variants, though these lacked detectable enzymatic activity. In the present study, we addressed these complementary limitations by developing an integrated approach that synergistically combines rational design principles with deep learning.

Specifically, we used rational design to define permissible regions within protein sequence space, which were subsequently subjected to BayesDesign optimization, resulting in the development of two libraries of NLuc variants (A and B). This hybrid methodology effectively harnesses the strengths of both approaches while mitigating their individual shortcomings. A schematic comparison of these methodological progressions is presented in Figure 1.

**Figure 1.**
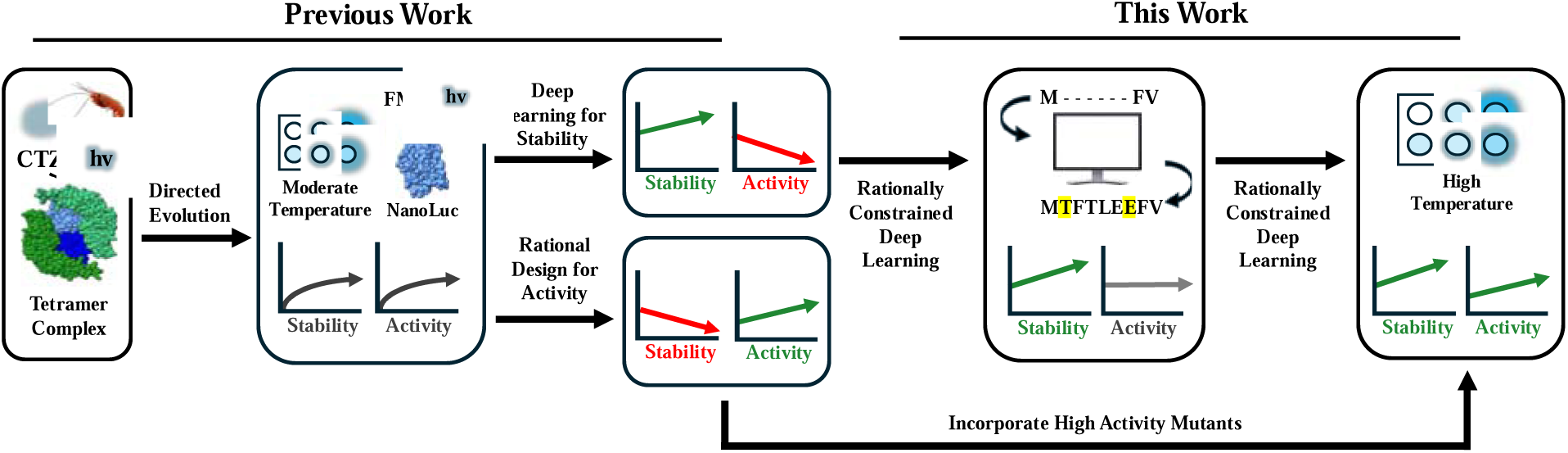
NanoLuc originated from deep sea shrimp via directed evolution. Subsequent efforts improved either stability using deep learning or activity through rational design. In this study, we integrate both approaches to jointly optimize stability and activity through two iterative rounds of computational prediction and experimental validation.

Figure 2 presents a comprehensive overview of five sequential rounds of NLuc engineering. While we previously reported thermostability – measured as percent solubility after heat treatment – for selected round 2 variants, we now provide complete data across all rounds for methodological comparison [21]. Initial rounds (1–3) primarily constrained the number of sites the deep learning model could mutate. Round 1 (1.01-1.10) was the least restrictive, requiring only 29 to 36 % homology with the original wild-type NLuc sequence. Round 2 was restricted to 48 to 96 % homology, and round 3 mutants were constrained to at least 99 % homology.

**Figure 2.**
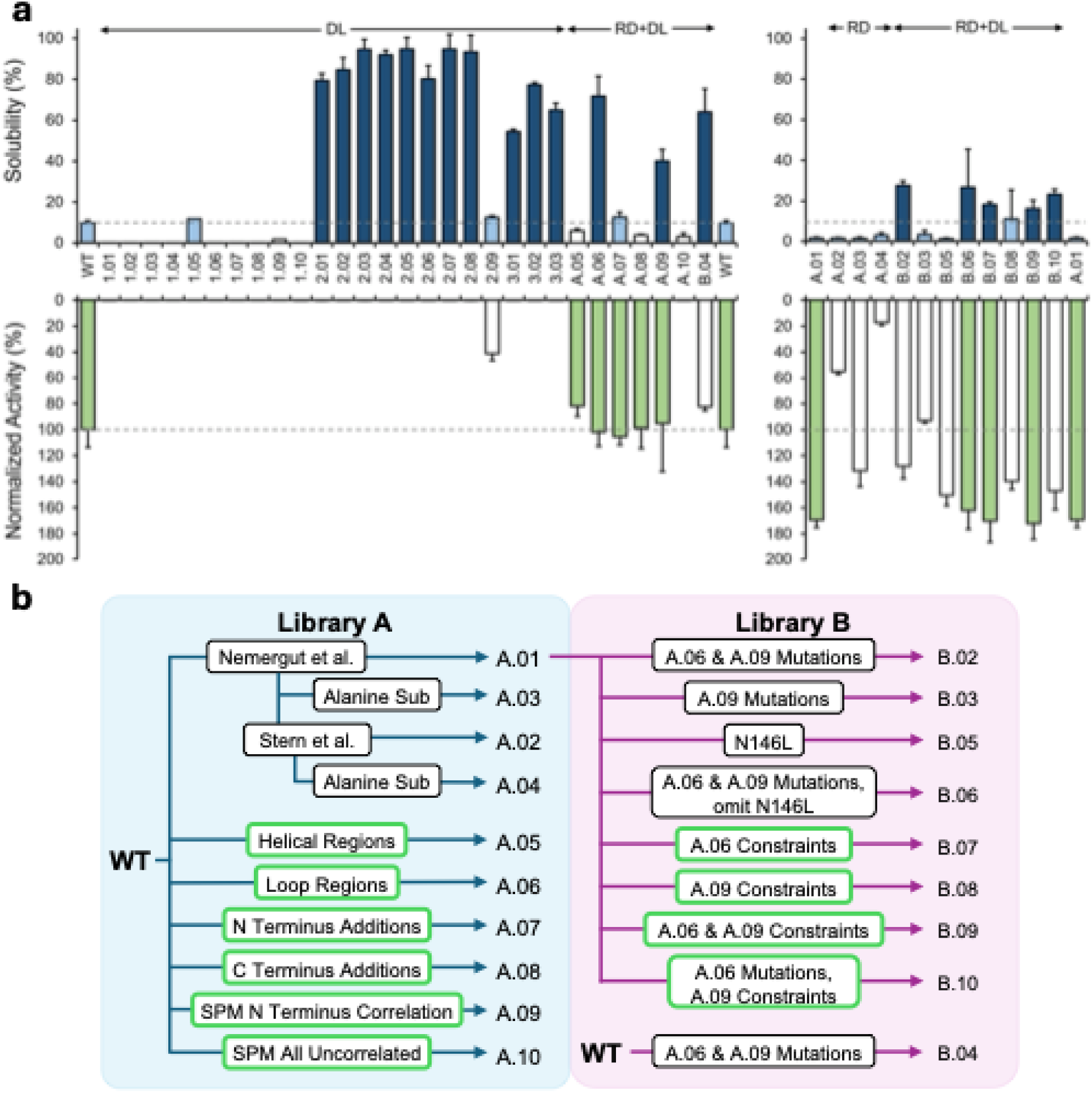
a) Comprehensive solubility and activity results from five sequential rounds of NLuc variants. Left chart presents variants derived from the wild-type sequence, while right chart presents variants derived from the A.01 sequence, with parent sequences shown on both sides of the charts for facile comparison. Solubility was measured at 57 °C (Rounds 1-3) or 60 °C (libraries A and B). Activity normalized to wild-type was measured at 30 °C (Rounds 1-3, library A) or 37 °C (library B). Dashed gray lines indicate baseline wild-type activity and solubility thresholds. Bar color indicates performance when compared to parent sequence, with white bars indicating lower performance, moderate shading indicating similar performance, and dark shading indicating greater performance. b) A schematic depicting the lineage (arrows from starting sequences) and design strategies (box texts) of variants in libraries A and B. Green borders represent mutations that were generated using BayesDesign.

**Figure 3.**
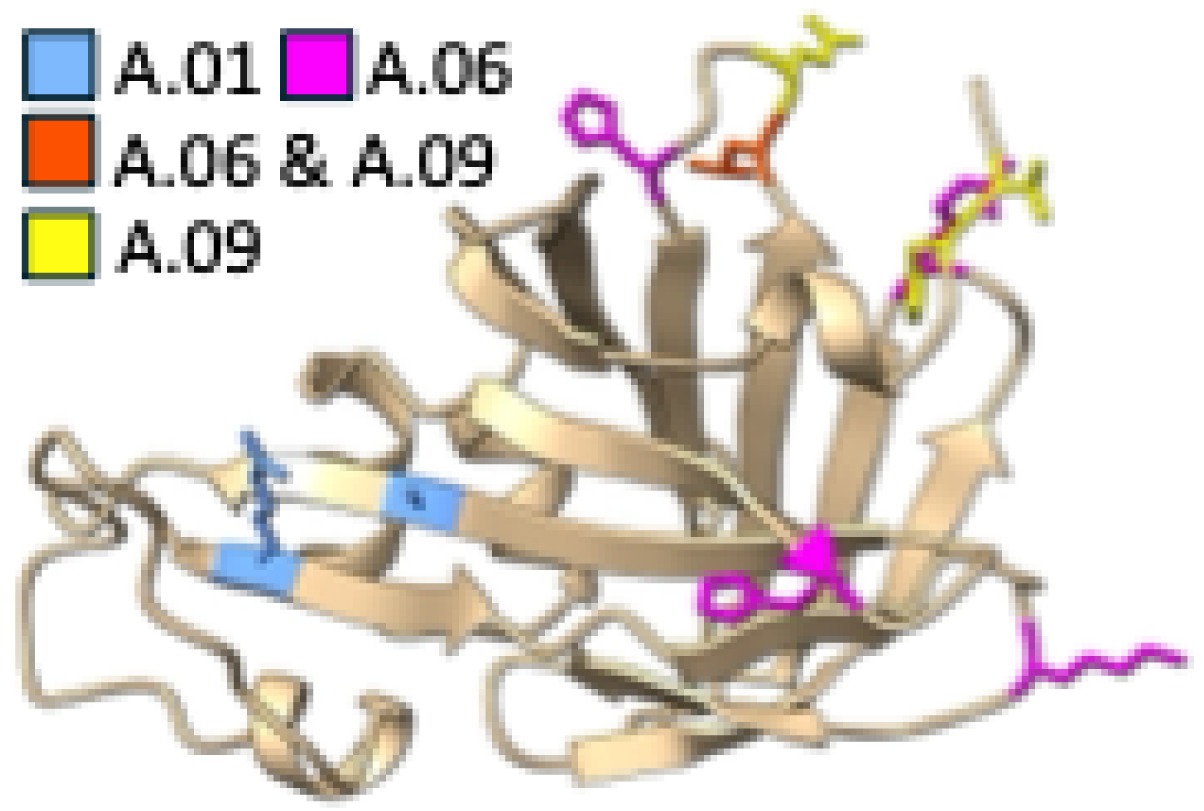
Structural and functional analysis of NLuc variants across libraries A and B. *Mapping of beneficial mutations from library A that were systematically recombined in library B, visualized on the NLuc crystal structure (PDB ID: 7SNT). Key mutation sites are color-coded according to their functional impact*.

Variants in rounds 2 and 3 showed markedly improved solubility at elevated temperatures, even when restricted to only two or three mutations. However, activity was consistently lost and typically undetectable. To overcome this limitation, our current approach combines rational design with BayesDesign to identify key regions and generate libraries A and B.

Library A consists of four sequences derived from work by Nemergut et al, and six sequences generated using a combination of rational design and BayesDesign. Variant A.01, a sequence previously characterized by Nemergut et al. as exhibiting enhanced activity with FMZ, was found to be highly active (169.4 ± 5.8 % of wild-type) but with compromised thermostability (1.5 ± 0.5 % of post-heat-treatment wild-type solubility). A.02 incorporates a stabilizing mutation identified in Round 3 (V40T) into the A.01 sequence, while A.03 and A.04 are alanine substitutions of residues 91 and 40, respectively. Variants 5-10 were generated by running the wild-type sequence through BayesDesign with a variety of masks. Each mask, which specifies which residue positions can mutate, was created using a unique rational design approach: only helix regions (A.05), only loop regions (A.06), C-terminus extension (A.07), N-terminus extension (A.08), areas dynamically correlated with the N-terminus (A.09), or residues with minimal dynamic correlation (A.10). Variant A.06 achieved remarkable stability enhancement (71.8 ± 9.6 % solubility compared to 9.7 ± 1.2 % for wild-type post-heat-treatment) while maintaining robust enzymatic activity (102.0 ± 10.9 %). Similarly, variant A.09 demonstrated substantial stability improvement (40.1 ± 5.6% solubility) while preserving 95.4 ± 36.9 % of wild-type activity. The alanine substitution at R91 (A.03) confirmed the importance of A.01 mutations for enhanced activity. The additional mutation in A.03 did not affect solubility but did decrease activity relative to A.01. Terminal extension variants showed that C-terminal modifications (A.07) modestly improved both stability and maintained activity, while N-terminal additions (A.08) decreased stability. Mutating residues with minimal dynamic correlation significantly decreased activity (A.10). Comprehensive numerical values for all variants’ activities and solubilities are provided in Table S1.

In library B, we systematically recombined mutations from the best-performing library A variants with all variants except B.04 using A.01 as the template sequence. B.04 was designed as a combination of A.06 and A.09 mutations. This combination did not provide synergistic stability effects since B.04 showed stability and activity below A.06, indicating that the A.06 and A.09 mutations are not synergistic. Generated variants B.02-03 and B.05-06 used combinations of mutations from A.01, A.06, and A.09. This recombination approach yielded diverse functional profiles, with B.02 resulting in a significant increase in stability but lower activity than A.01. B.06 demonstrated improvements in stability and retained the activity of A.01 albeit with large stability error bars due to poor expression (see Figure S1). Additionally, we applied a second round of guided deep learning to combinations of A.01, A.06, and A.09 mutations, generating variants B.07-B.10. This approach yielded several exceptional variants. B.09 achieved the highest activity in our dataset (172.1 ± 12.3 % of wild-type) and maintained improved stability (16.1 ± 4.3 % solubility). Variant B.07 closely matched this activity level (170.0 ± 16.4 %) with slightly higher stability (18.0 ± 1.5 % solubility).

Overall, six of the nine library B variants exhibited substantial enhancements in stability compared with A.01, while simultaneously maintaining significantly increased enzymatic activity relative to wild-type NLuc. It should be noted that replacing the homology-based approach from the first three rounds with this rational design-based approach produced many variants with maintained activity. While previous rounds successfully produced soluble enzymes up to 95 °C, the enzymes lacked detectable activity. Results for libraries A and B indicate that targeting disordered regions (defined here as regions that are not an alpha helix or beta sheet) distant from the active site and enhancing dynamically coupled networks were particularly effective approaches for balancing stability and function. These findings highlight the complex nature of the stability-activity trade-off and how a modified approach combining both rational design and deep learning can be used to mitigate it.

### Computational investigation of NLuc variants

Figure 4a illustrates the comprehensive mutational sequence space explored across libraries A and B, revealing distinct patterns in BayesDesign’s sampling strategy. Across most permissible positions, BayesDesign predominantly sampled only two amino acid variations, suggesting a focused convergence toward specific biochemical properties at these sites. This limited mutational diversity likely reflects the algorithm’s targeting of evolutionarily conserved positions, where only a narrow range of substitutions can occur without compromising structural integrity. Notably, variant B.09 stands as an exception to this pattern, exhibiting unique mutations at positions V2H and K125Q that were not observed in any other variant. These exclusive substitutions in B.09 – replacing a hydrophobic valine with a positively charged histidine at position 2 and substituting a positively charged lysine with a polar glutamine at position 125 – suggest the algorithm explored alternative electrostatic and hydrogen-bonding networks in this design iteration. The restricted mutational diversity at most positions, contrasted with these unique variations in B.09, highlights the delicate balance between conservation and innovation in our protein engineering approach, where BayesDesign effectively navigated the vast theoretical sequence space to identify viable candidates with enhanced functional properties.

**Figure 4.**
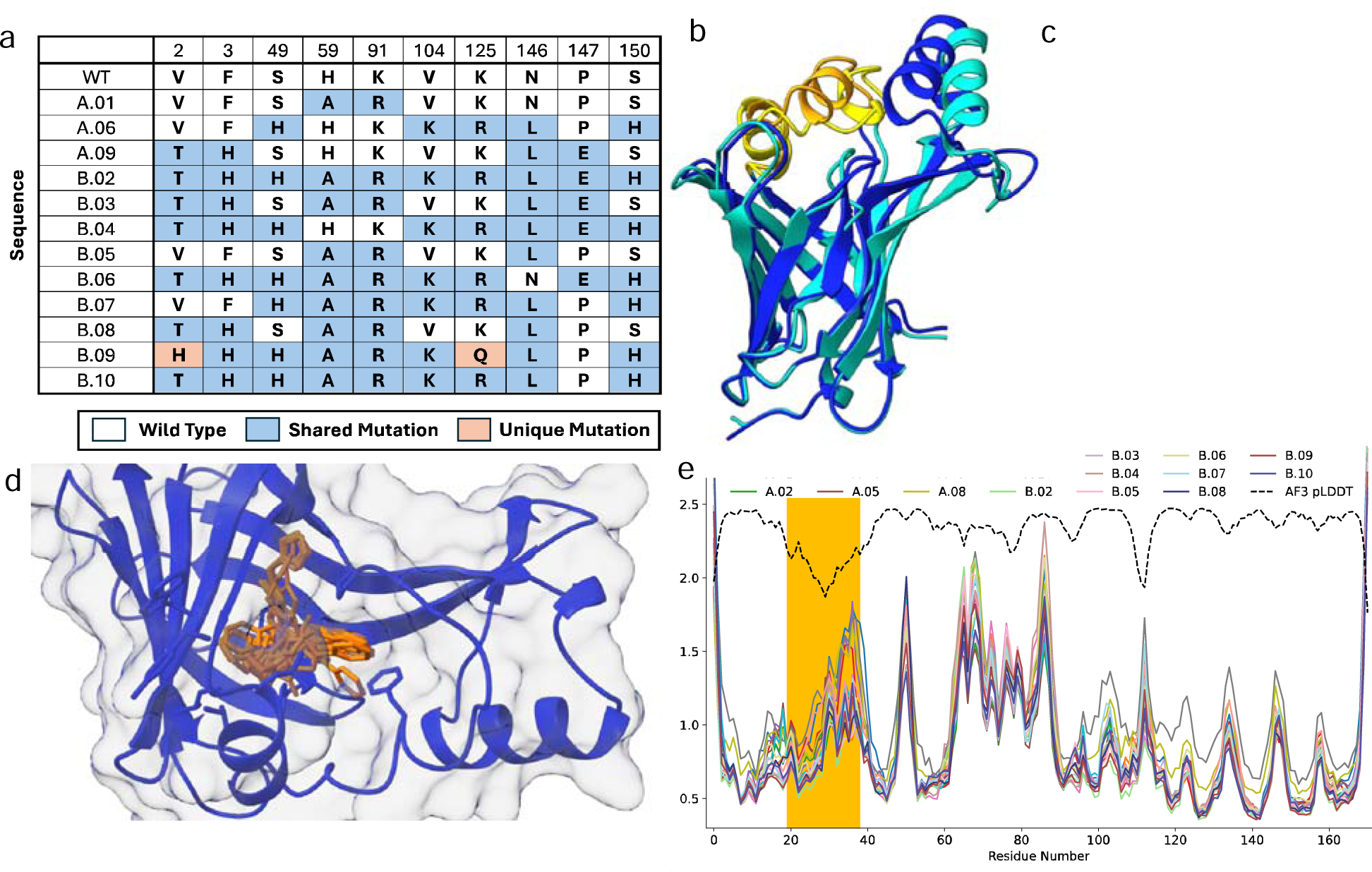
Computational Analysis of mutants. a) Sequences of the most important mutants investigated. Top row indicates the residue position; letters stand for amino acid codes. b) The side and c) top views of two conformations assumed of all mutants when folded repeatedly with AF3 in absence of a substrate. Golden helix indicates area of high uncertainty in AF3 pLDDT scores. d) Predicted binding position of all FMZ substrates in one example backbone. e) C-alpha RMSF of all variants derived from MD simulations. The black dashed line shows a rescaled (divide by 100 add 1.5) pLDDT average score across all variants from AF3. The yellow highlight corresponds to the possible lid region highlighted in b) and c).

Structural ensemble analyses are presented in Figure 4b (side view) and Figure 4c (top view), revealing distinct conformational states sampled by AF3 [30]. Remarkably, each mutant sequence consistently yielded two alternative conformational predictions with high confidence. As these structural models were generated in the absence of substrate, the dual conformations likely represent functionally relevant states that correspond to “open” and “closed” conformations of the enzyme. Of particular significance is the orange-coded loop region spanning residues 19-31, which contains transient helical motifs and exhibits the highest uncertainty in pLDDT scores across all AF3 predictions (see also Figure S2). This region of heightened conformational ambiguity appears to constitute a flexible lid domain that may regulate substrate access to NLuc’s beta-barrel active site. The conformational heterogeneity captured in these models aligns with established principles of enzyme dynamics, wherein flexible elements often modulate catalytic function through controlled transitions between discrete structural states. This computational evidence for conformational plasticity provides valuable insights into the structural basis of NLuc’s catalytic mechanism and offers a framework for understanding how mutations might influence the dynamic equilibrium between functional states.

Figure 4d provides a detailed examination of the binding site architecture across NLuc variants. We employed AF3 to model all mutant sequences in complex with the substrate FMZ, enabling assessment of potential changes in substrate-binding interactions. While conformational heterogeneity was observed comparable to that seen in apo-protein predictions, the location of the binding pocket remained remarkably consistent across all variants. Notably, FMZ possesses ten energetically stable conformers, and AF3 predictions sampled a diverse spectrum of these rotameric states within the binding site [31]. Given the recognized limitations of AF3 in accurately predicting fine-grained ligand-protein interactions, particularly for flexible small molecules, we emphasize the qualitative consistency in substrate docking behavior rather than specific interaction energetics. The preservation of binding site topology across variants suggests that the mutations introduced by BayesDesign primarily modulate protein stability and dynamics without fundamentally altering the substrate recognition mechanism. The subtle structural adaptations that likely underlie the observed enhancements in catalytic efficiency appear to be below the resolution threshold detectable by comparative structural modeling.

Figure 4e illustrates the dynamic behavior of all 20 NLuc variants through 15 independent 10 ns molecular dynamics simulations each, with FMZ in the binding site (see also Figure S3). Throughout all simulations, FMZ remained stably bound in the active site without requiring additional constraints, validating the appropriate electrostatic environment for maintaining the non-covalent holo-conformation. Interestingly, the alpha-helical lid region exhibited reduced conformational fluctuations compared with what might be expected from the conformational heterogeneity observed in Figures 4b and 4c. This discrepancy likely stems from the presence of FMZ in the binding site, which appears to stabilize the “open” conformation and restricts the conformational sampling of the lid region during simulation timescales. We overlay in Figure 4e (dashed black line) the average pLDDT confidence scores from AF3 predictions across all mutants with bound FMZ, revealing a significant inverse correlation with the root-mean-square fluctuation (RMSF) values from MD simulations (Pearson correlation of -0.5, p < 0.001). This correlation suggests that regions with lower predicted structural confidence tend to exhibit greater dynamic flexibility. A notable exception occurs in the segment spanning residues 60-90, which displays elevated RMSF values despite relatively high pLDDT confidence scores. This region corresponds to a segment that was poorly resolved in the crystallographic structure (PDB ID: 7SNT), with electron density missing for 11 residues. Interestingly, comparing the RMSF between AF3 generated structures for all mutants with the average RMSF across all simulations, resulted in nearly identical RMSF spectra, only modified by an amplitude of 2. This suggests that in some cases AF3-created structural ensembles may contain similar information to molecular dynamic simulations (see Figure S4).

### Detailed Biophysical Characterization of Selected NLuc Variants

To comprehensively evaluate the most promising NLuc variants, we conducted in-depth analyses of thermostability, pH dependence, and spectral properties using the three most interesting sequences that resulted from the expert guided deep learning approach: A.06, B.07, and B.09 and compared them with the wild-type and A.01 sequences. For reference, decay trends as well as kinetic parameters for each variant are reported in Figures S8 and S9, with thermostability data reported below.

### Thermostability Profiles

Temperature-dependent solubility measurements revealed significant stability improvements between the original and designed variants (Figures 5a and 5b, Table S2). We calculated the temperature at which 50 % solubility is maintained (T) as a quantitative measure of thermostability (Table S3). The rationally designed variant A.06 exhibited exceptional thermostability with a T of approximately 64 °C, representing about 9 °C increase compared with wild-type (T = 55 °C). This substantial improvement confirms the effectiveness of targeting disordered regions distant from the active site. In contrast, the high-activity variant A.01 showed reduced thermal stability (T = 51 °C), consistent with its compromised solubility at elevated temperatures reported in Figure 2.

**Figure 5.**
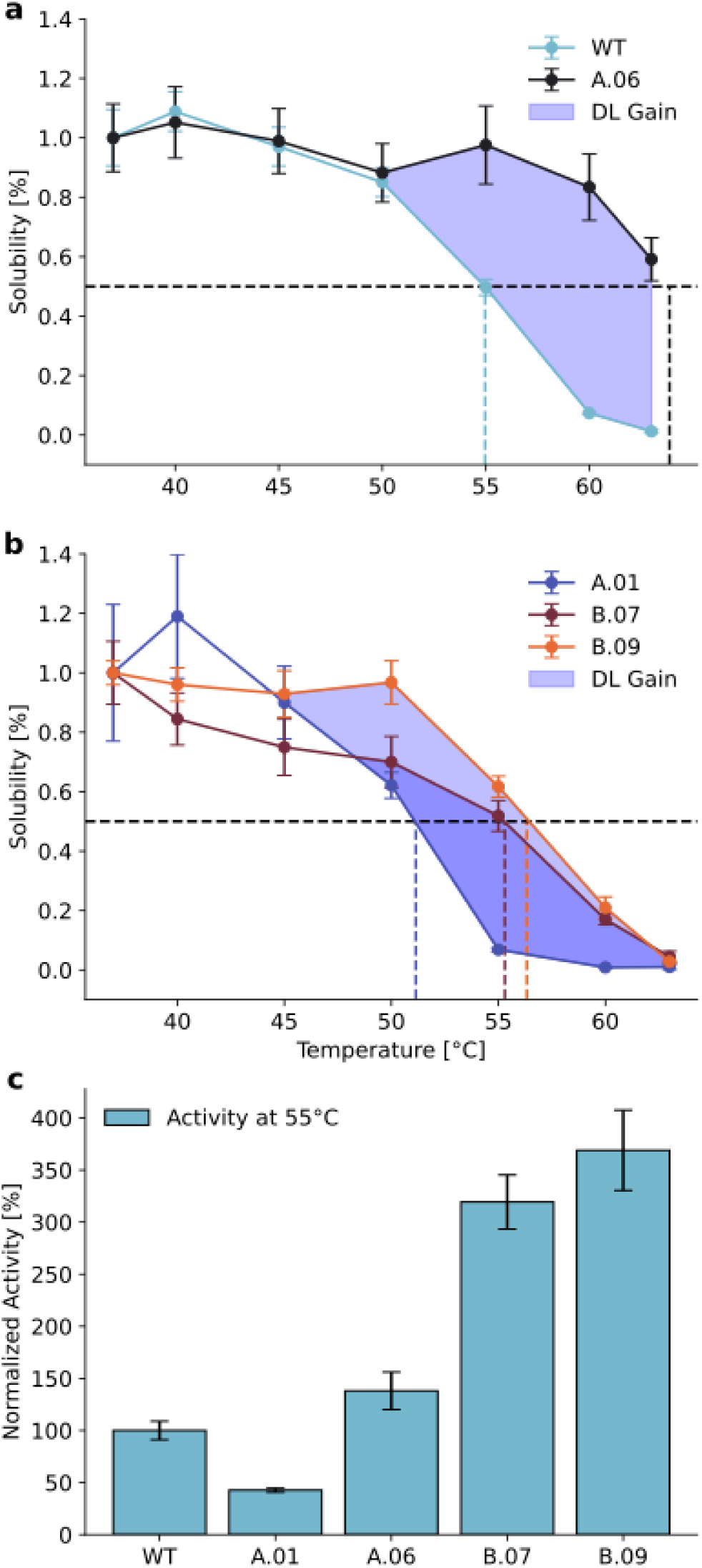
Detailed experimental analysis of promising NLuc variants. a) Solubility of variant A.06 compared with wild-type. Blue shaded area indicates solubility gains achieved through our deep learning (DL) protocol. b) Solubility comparison of A.01-based variants. Blue shading highlights improvements attributable to DL methodology. Dashed lines identify T50 values. c) Activity of NLuc variants at elevated temperature, normalized to wild-type.

**Figure 6.**
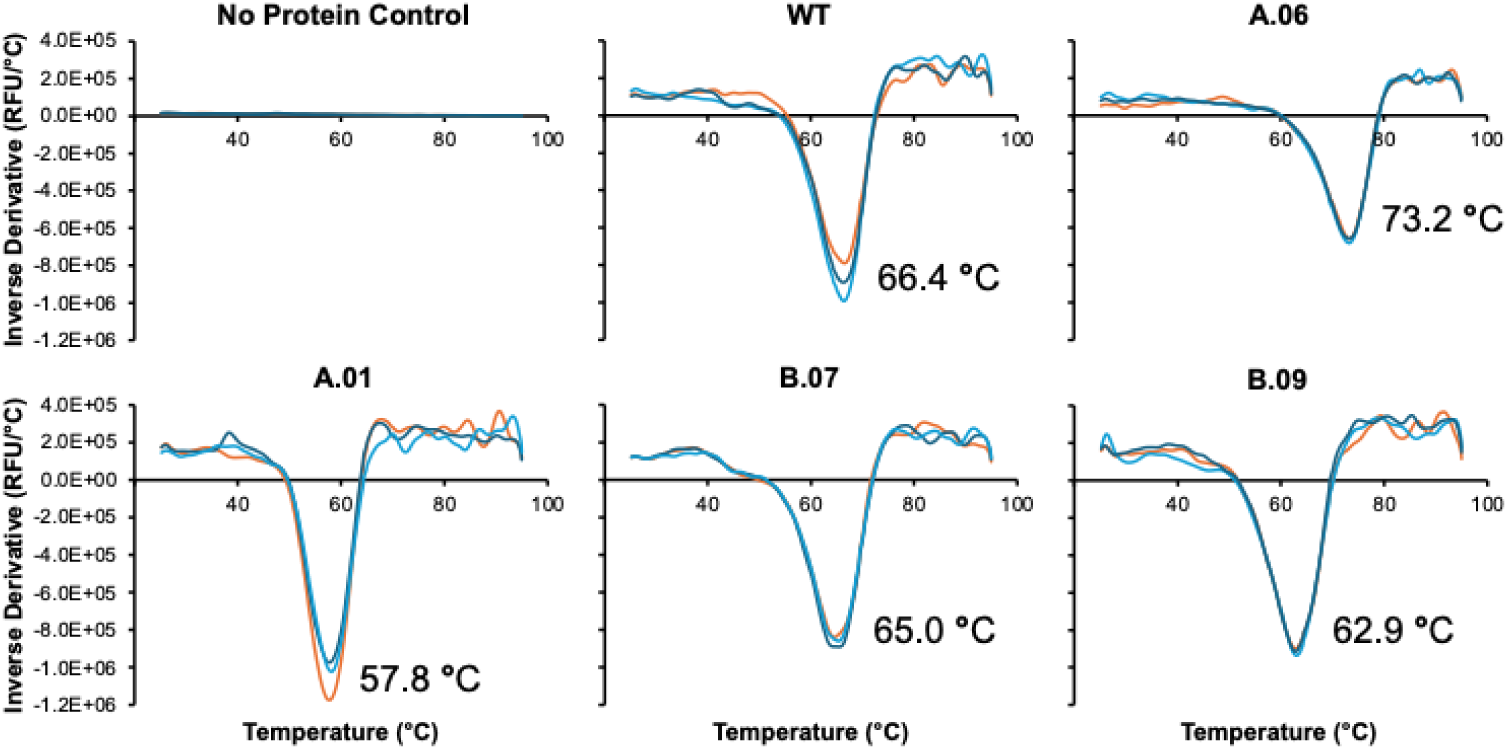
Thermal shift assay profiles of selected variants along with No Protein Control samples. The minimum of the inverse derivative of the fluorescent profiles represents the melting point of the variant. Each line represents one replicate where each variant was sampled in triplicate.

Notably, our hybrid designs B.07 and B.09 successfully preserved the enhanced activity of A.01 with significantly higher thermostability than A.01, with T values of 55 °C and 56 °C, respectively. When examining the complete thermal denaturation curves reported in Figure 5, A.06 maintained over 83 % solubility at 60 °C and retained substantial solubility (59 %) even at 63 °C. In contrast, wild-type NLuc retained only 7.4 % solubility at 60 °C, while A.01 was almost completely insoluble (0.9 %) at this temperature. The engineered variants B.07 and B.09 showed intermediate stability, retaining 17 % and 21 % solubility at 60 °C, respectively.

To validate the solubility data, we conducted thermal shift assays to determine the melting temperatures of the variants, see Figure 6. The melting temperature is defined as the minimum of the inverse derivative of the fluorescence profile for a given sample. The measured melting temperatures followed the same relative ranking as the solubility assays, where variant A.06 exhibited a 6.8 °C increase in comparison to the wild-type sequence and variants B.07 and B.09 exhibited increases of 7.2 and 5.1 °C in comparison to A.01, respectively.

### Spectral Characteristics with Alternative Substrates

We examined the spectral properties of the selected variants with two distinct substrates – FMZ, (the primary substrate for NLuc) and CTZ (an industrially beneficial alternative substrate) – across the wavelength range of 400-600 nm (Figure S5, Table S4). With FMZ, wild-type NLuc exhibited a characteristic emission peak at approximately 460 nm. The high-activity variant A.01 maintained a similar spectral profile. Variant A.06 maintained the native emission spectrum shape.

The hybrid variants B.07 and B.09 showed no significant shifts in peak wavelength, suggesting that the mutations primarily affect quantum yield rather than the electronic state of the excited chromophore.

With CTZ as substrate, all variants maintained the characteristic, red-shifted emission profile compared with FMZ, with peak emission at approximately 480 nm (Table S5 and Figure S6). A.01 was previously engineered for CTZ activity, and the hybrid variants retained that beneficial trait [29]. Indeed, A.01, B.07, and B.09 were over five times as active as wild-type and A.06 with the CTZ substrate (Figure S6).

### pH-Dependent Activity Profiles

We characterized the pH dependence of enzymatic activity across pH 4-9 to assess catalytic robustness under varying conditions (see Figure S5, Table S6). All variants displayed a pH-activity profile typical of luciferases, with activity increasing from acidic to neutral/basic conditions. Wild-type NLuc showed optimal activity at higher pH, with substantially reduced activity below pH 6. The high-activity variant A.01 maintained elevated activity across the pH range and retained significant activity at pH 5 (9.8 × 10 RLU), representing a 5.3-fold increase over wild-type at this pH.

Interestingly, variant A.06 exhibited similar profile to the wild-type sequence verifying that mutations which improve thermostability do not decrease pH stability. The hybrid variants B.07 and B.09 maintained both enhanced activity and pH tolerance inherited from A.01. B.09 achieved the highest overall activity, representing a 22 % increase over wild-type at pH 9. Notably, all engineered variants showed minimal activity at pH 4, indicating that extreme acidic conditions remain challenging for NLuc function regardless of the introduced mutations.

### Enhanced Catalytic Activity at Elevated Temperatures

Temperature-dependent activity assays revealed striking differences in the thermal performance profiles of NLuc variants. At 55 °C, variant A.01 exhibited significantly diminished activity (42.7 ± 1.9% of wild-type), consistent with its reduced thermostability and accelerated denaturation kinetics (Figure 5). In contrast, the stabilized A.06 variant demonstrated enhanced catalytic efficiency (138.1 ± 18.0 % relative to wild-type) under these conditions. Most remarkably, the engineered variants B.07 and B.09 displayed exceptional thermal activation, achieving 319.3 ± 26.0 % and 369.0 ± 38.6 % of wild-type activity, respectively (Figure 5). This substantial performance enhancement at elevated temperatures highlights the potential application of these variants in high-temperature sensing and industrial bioprocesses, where thermostable biocatalysts with enhanced catalytic efficiency could be beneficial [18–20].

Collectively, these comprehensive analyses demonstrate that our integrated rational design and deep learning approach successfully generated NLuc variants with enhanced thermostability that leads to improved catalytic properties at elevated temperatures and maintains pH tolerance and expanded substrate compatibility. The exceptional performance of variants B.07 and B.09 highlights the effectiveness of our hybrid engineering strategy in addressing the historically challenging trade-off between protein stability and activity.

## Conclusion

Our study overcomes the stability-activity trade-off in NLuc luciferase engineering through an integrated rational design and deep learning approach. By targeting disordered regions distant from the active site while preserving dynamically coupled networks, we developed variants with significantly improved thermostability (T increases up to 5.19 °C) and enhanced activity at elevated temperatures (up to 369% of wild-type at 55 °C). These engineered luciferases maintain wild-type NLuc’s pH tolerance with optimal function at biological pH and improved compatibility with alternative substrates. Our hybrid methodology provides a generalizable framework for engineering enzymes with limited sequence homology, offering enhanced tools for bioluminescence imaging, biosensing, and protein interaction studies.

## Methods

### Generation of NLuc variants

We followed the procedure outlined in Figure 7. NLuc variants were designed using BayesDesign, a deterministic deep learning-based algorithm that predicts the most probable sequence given structural and amino acid constraints. Structural constraints were defined by the Cα positions of wild-type NLuc, as specified in PDB ID: 7SNT, while amino acid constraints were based on the wild-type NLuc sequence, see Table S7 for all sequences. For each prediction, a subset of residues was selected for optimization, typically within flexible regions such as residues X–Y in a loop. BayesDesign was implemented via an established Google Colab notebook using sequence mask regions and input PDB structures as constraints [21]. All sequences were generated using the PDB structure of NLuc or its variants, primarily PDB ID: 7SNT [32], except for shortest path analyses – described below –, which used PDB ID: 5IBO [33]. Structures were obtained from the Protein Data Bank. Sequence generation followed a greedy decoding algorithm, proceeding from the N-terminus to the C-terminus. Positions designated as “fixed positions” remained unchanged during sequence optimization, ensuring the preservation of critical structural or functional elements.

**Figure 7.**
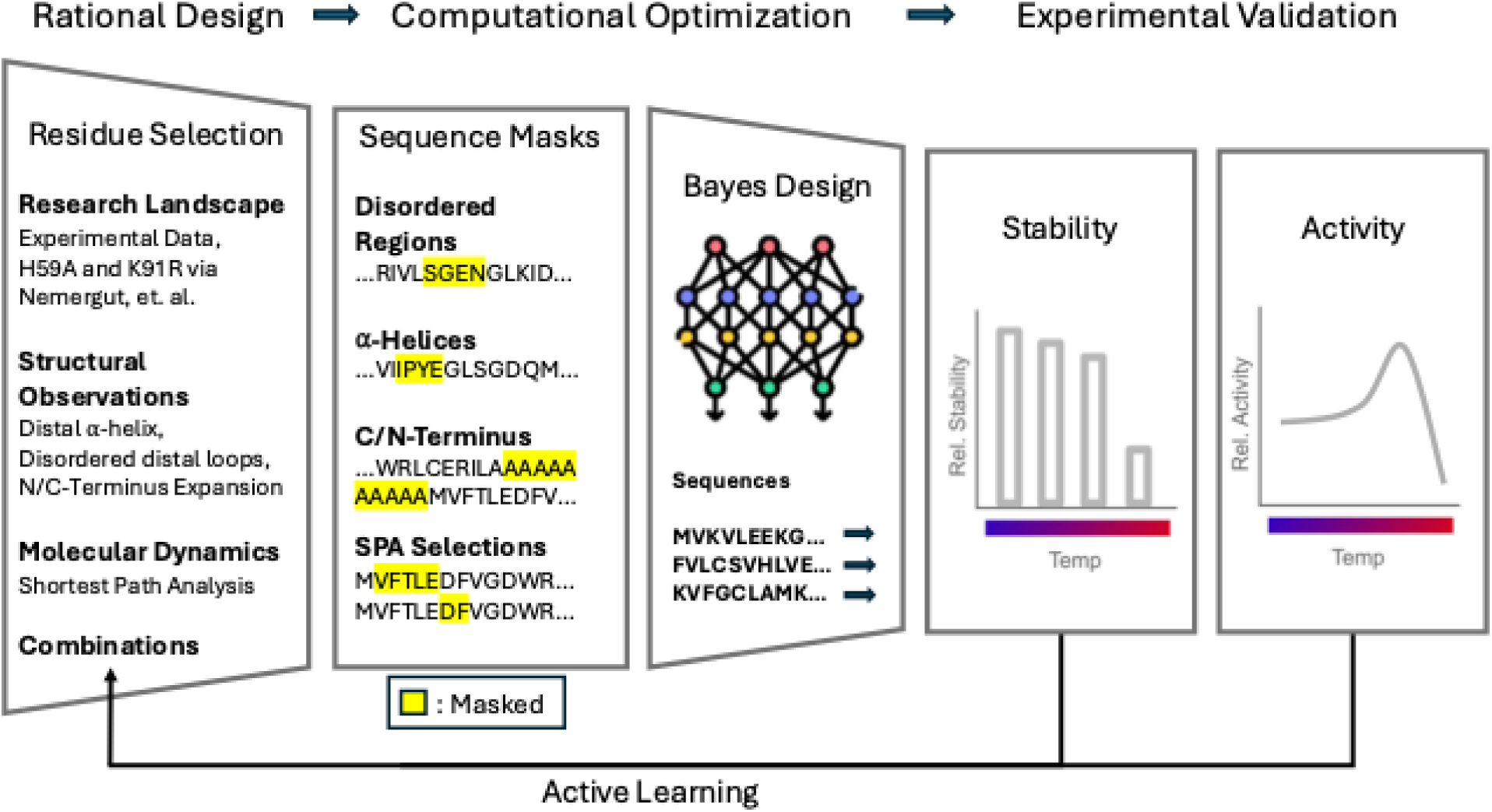
Integrated rational design and deep learning pipeline for enzyme engineering. Structure-guided analysis of literature and molecular simulations identifies key sequence regions for targeted optimization. The BayesDesign algorithm iteratively refines these sequences to discover variants with enhanced properties. Experimental validation provides feedback for subsequent rounds of expert-guided residue selection, creating an active learning cycle that efficiently navigates the sequence-function landscape.

### Generation of variant library A

Four sequences were selected based on combinations of mutations identified in previous studies: H59A and K91R from Nemergut et al. [29] and V40T from Stern et al. [21]. The amino acid sequence numbering in this work is consistent with the original sequence reported by Hall et al. [16]. Sequence combinations are as follows: A.01 (H59A, K91R), and A.02 (V40T, H59A, K91R). Sequences A.03 and A.04 were constructed using Alanine substitutions in place of mutations identified in A.01 and A.02, respectively, making combinations: A.03 (H59A, K91A), and A.04 (V40A, H59A, K91A). Additionally, six sequences were designed by generating NLuc variants with BayesDesign, which entailed generating a mask (a list of residue positions that BayesDesign was allowed to mutate) and applying said mask to the wild-type sequence. These masks were created using a variety of rational design techniques, namely: helix regions (residues 62–65 and 71–84), loop regions (49–52, 101–106, 121–125, and 145–150), N-terminal additions (−4-0), C-terminal additions (172-176), dynamic correlations with the N-terminus (2-6, 146-147), and residues with minimal dynamic correlation (7, 8, 21, 27, 51, 63-64, 85-86, 99, 106, 120, 128, 145, 149). Dynamic correlations were identified via shortest path analysis of MD simulations of the wild-type enzyme. Structures submitted to BayesDesign for terminal modifications were generated by appending alanine residues to the wild-type 7SNT structure in Chimera [34]. For mutation details, see Table S8.

### Shortest Path Analysis

Analysis of molecular dynamics simulations from our previous work was conducted to inform the selection of mutation sites [21]. The complex was prepared by docking FMZ with AutoDock Vina v1.2.3 to NLuc (5IBO) [35]. Parameterization of FMZ used OpenFF’s SMIRNOFF force field [36] while Amber ff14SB force field was used for the protein [37]. Following solvation with explicit TIP3P water, charge neutralization, equilibration, minimization, and NPT equilibration, three independent 100 ns simulations were performed. The above processes were conducted using OpenMM’s molecular dynamics engine [38]. All residues that came within 3 angstroms of the ligand across all MD simulations, as identified by ChimeraX 1.4, were considered active site residues [36]. The resulting trajectories were analyzed using the web server developed by Casadevall et al. [39], which performs shortest path method (SPM) analysis. SPM analysis is an algorithm that uses MD trajectories to determine amino acids that move in concert as a measure of interacting residues. Two SPM analyses were run, the first used default constraints (significance threshold 0.3, distance threshold 6) where residues selected for mutation were determined from locations excluded from the active site that showed similar movement. This defined the mask for variant A.09. The second analysis used maximally permissive constraints (significance threshold 0.1, distance threshold 10.). The mask for variant A.10 was created by selecting residues that were not identified by the second analysis and were not part of the active site.

### Generation of variant library B

Library B was derived from the best-performing sequences in library A, combined either directly (by maintaining mutations) or dynamically (by using library A mutations as mutable positions in BayesDesign with other positions constrained). Four direct combinations were generated as follows: B.02 (A.01, A.06, A.09), B.03 (A.01, A.09), B.04 (A.06, A.09), and B.05 (A.01, N146L). Sequence B.06 incorporated mutations from A.01, A.06, and A.09, except N146L, which was reverted to wild-type asparagine.

Four dynamic combinations were generated using a modified 7SNT structure as a backbone, integrating sequences from library A into the wild-type structure using the ChimeraX rotamer tool [40–42]. These structures were submitted to BayesDesign with mutation positions defined as follows: B.07 (backbone: A.01, mutable positions: A.06), B.08 (backbone: A.01, mutable positions: A.09), B.09 (backbone: A.01, mutable positions: A.06, A.09), and B.10 (backbone: A.01, A.06, mutable positions: A.09). For mutation details see Table S8.

### Structure prediction

Protein structures were predicted using AF3 [30]. Model weights for self-hosted AF3 installation were made available by DeepMind upon our request – these are needed to fold with arbitrary substrates, like FMZ or CTZ. Each sequence in libraries A and B were input into a *json* file along with random seeds and processed by AF3. The random seeds allowed for analysis of the variations in conformation seen in Figures 4, S2 and S7. Each sequence produced five outputs ranked based on the highest average pLDDT confidence score. The output with the highest pLDDT score was used. Structures with the substrate FMZ were created by including the substrate in the same input file as the sequence. The output files from AF3 contained a pLDDT value for each atom within a structure. Here, the pLDDT values for each α-carbon were used.

ChimeraX 1.8 and 1.9 were used to visualize the AF3 outputs [40–42]. pLDDT values were displayed throughout the structure with the AlphaFold Error Plot tool on ChimeraX. The Matchmaker tool was used in ChimeraX to align protein structures to compare sequences and substrate binding. To understand how the protein was interacting with the substrate, the Contact tool found in ChimeraX 1.9 was applied to the protein-substrate complex.

### Plotting and visualization

Visualization and analysis were performed using Microsoft Excel, Matplotlib v3.7.1 [43] in Python v3.10, and UCSF ChimeraX 1.5 [40–42]. Experimental data in Figure 2a was visualized using Excel and flow charts from Figure 2b in PowerPoint. Comparative analyses in Figures 3, 4e, and 5 were generated using Matplotlib. Structural visualizations in Figures 3, 4b, and 4d were rendered in ChimeraX using AF3-predicted protein models. Mutational effects in Figure 3 were modeled using the rotamer optimization tool in ChimeraX with Dunbrack 2010 rotamer libraries [44].

### Protein Flexibility Analysis

Molecular dynamics (MD) simulations were performed using GROMACS 2024.3 with the CHARMM36m force field (July 2022 release) [45–53]. AF3-predicted structures of NLuc variants served as initial conformations. Substrate parametrization for FMZ was conducted using the CHARMM General Force Field (CGenFF v4.6) via the ParamChem web server [54], with partial charge assignments and dihedral parameters optimized according to established protocols. Ligand molecular structures were prepared and optimized using Open Babel v3.1.1[55, 56] with the MMFF94s force field prior to parametrization.

Each protein-ligand complex was centered in a dodecahedral simulation box with a minimum distance of 10 Å between the complex and box edges, then solvated with CHARMM-modified TIP3P water molecules [30]. Systems were prepared with standard protonation states (N-terminus as NH, C-terminus as COO) and neutralized by adding Na counterions to achieve electroneutrality. Energy minimization was performed using the steepest descent algorithm with a 0.01 nm step size and convergence criterion of maximum force below 1000 kJ mol ¹ nm ¹.

System equilibration consisted of sequential NVT and NPT ensemble simulations. The NVT step maintained 300 K for 100 ps using the velocity rescaling thermostat (τ = 0.1 ps), followed by 100 ps NPT equilibration at 1 bar pressure controlled by the Berendsen barostat with stochastic cell rescaling (τ = 2.0 ps). Both steps used a 2 fs integration time step with the leap-frog algorithm.

Long-range electrostatics were treated with the particle mesh Ewald (PME) method using a 1.2 nm real-space cutoff. Van der Waals interactions were truncated at 1.2 nm with a smooth switching function applied from 1.0 nm. All bond lengths involving hydrogen atoms were constrained using the LINCS algorithm (4th order expansion with 1 iteration). The substrate was constrained with 1000 kJ mol ¹ nm ².

Production simulations were conducted for each of the 20 variants (wild-type and 19 engineered variants). For statistical robustness, we performed 7 independent 100 ns simulations per variant using different initial velocity distributions, yielding a cumulative 14 μs of simulation data. All production runs maintained NPT conditions (300 K, 1 bar) with identical parameters as the equilibration phase. Trajectories were saved every 10 ps for analysis.

Root mean square fluctuation (RMSF) values for Cα atoms were calculated using the GROMACS `gmx rmsf` module after least-squares fitting to the initial structure to remove translational and rotational motion. For each variant, RMSF profiles were averaged across all 15 independent simulations to ensure statistical significance and minimize bias from any single trajectory. These averaged RMSF values were then mapped onto the protein structures and correlated with experimental activity and stability measurements.

### Expression of NLuc Variants

Cell extract for protein synthesis was prepared from *E. coli* BL21-Star™ DE3 cells (Invitrogen, Carlsbad, CA, USA) in 2xYT media. Cell-free protein synthesis (CFPS) was performed as reported previously [21, 57]. 1 L cultures were induced with isopropyl β-d-1-thiogalactopyranoside IPTG at an OD600 of 0.5 to 0.7 and harvested at an OD600 of 2 to 4.

Cells were washed and then lysed with an Avestin Emulsiflex B-15 homogenizer (Avestin, Ottawa, Canada) at 21000 psi over 3 system passes. The lysate was then centrifuged for 30 minutes at 12000 RCF. The supernatant was collected and then used in CFPS reactions at 25% (v/v). PANOx-SP prepared as reported previously was added to CFPS reactions [58]. The DNA template was added to CFPS reactions as unpurified PCR product at 33% (v/v). Protein yield was determined by drying 2 μL of unpurified CFPS product on filter paper, washing the sample 3 times with TCA (trichloroacetic acid), and measuring the percentage of incorporated 14C-leucine (added at 5 μM concentration in the initial CFPS reaction preparation) using a liquid scintillation counter [59].

### Activity and stability assays

NLuc mutants were expressed using CFPS for 3 h at 37 °C and 280 rpm. Solubility assessment was performed as previously reported [21]. Following synthesis, each reaction was sampled for scintillation counting and then divided into 10-12 µL aliquots. Each aliquot was heat treated within the range of 37 °C to 95 °C for 15 min in a PCR thermal cycler followed by centrifugation for 15 min at 16100 g and 4 °C. The supernatant was sampled in triplicate for scintillation counting to determine solubility. All activity assays were performed in a 96-well plate and measured in a Biotek (Winooski, VT, USA) SynergyMx plate reader. Enzyme activity was determined by first diluting the protein 500x in 1x PBS buffer and then adding 2 μL of the protein dilution to 75 μL of assay reagents per well, which includes 34.7 μL 1x PBS, 39.5 μL Nano-Glo® Luciferase Assay Buffer, and 0.79 μL Nano-Glo® Luciferase Assay Substrate (Promega, Madison, Wisconsin, USA). Activity assays for rounds 1-3 and library A were performed at 30 °C and measured after 3 min while activity for library B was run at 37 °C. Protein sequences from Figure 5 were heat treated at 55 °C for 15 min, followed by a subsequent freeze/thaw cycle for one day, and then measured for activity at 55 °C after 3 min. Activity was normalized to the total protein yield. Each round was normalized to the WT protein at their respective temperatures. CTZ sensitivity was determined by diluting protein 500x in 1x Dulbecco’s PBS. 2 μL of diluted protein was added to 75 μL of assay reagents, which include 34.7 μL 1x Dulbecco’s PBS, 39.5 μL Gaussia Glow Assay Buffer, and 0.79 μL 100x Coelenterazine (Pierce™ Gaussia Luciferase Glow Assay Kit, Thermo Scientific, Waltham, MA, USA).

Thermal shift assays were performed as reported previously [60]. Proteins were first purified using the Strep-Tactin®XT 4Flow® high-capacity Spin Column Kit (IBA Life Sciences, Gottingen, Germany) as specified by the manufacturer. The columns were washed 2 times with Buffer W, before eluting to the specified high concentration option. Liquid scintillation counting was used to determine purified protein yields. Melting temperature was determined using the Protein Thermal Shift™ Dye Kit (Thermo Fisher Scientific, Carlsbad, CA). A hydrophobic dye is used that fluoresces when bound to hydrophobic regions of a protein upon denaturation. The thermal shift reactions contained the following composition: 5 μL Protein Thermal Shift Buffer, 2.5 μL Diluted Protein Thermal Shift Dye (8×) diluted with water, and 12.5 μL Strep-Tactin®XT Elution Buffer (Buffer BXT) containing 500 ng of purified protein sample. Reactions were performed in triplicate in a FrameStar® 96 Well Semi-Skirted PCR Fast Plate (Midsci, St. Louis, MO) and covered with a MicroAmp® Optical Adhesive Film (Applied Biosystems, Thermo Fisher Scientific). The assay was run in a StepOnePlus Real-Time PCR System (Applied Biosystems) at standard ramp speed covering a range of 25 to 95 °C.

### pH Assay

The pH assay buffers were prepared according to previously reported protocols [16], where buffers contained 100 mM citrate, HEPES, or tricine, 1 mM DTT, 10 mM MgSO4, 0.5% Tergitol®, and 0.05% (v/v) Antifoam 204, titrated to their respective pH values. For the pH assay, proteins were diluted 500x into their respective pH buffers, then added to 75 μL of assay reagents per well, which includes 74.21 μL assay buffer and 0.79 μL of Nano-Glo® Luciferase Assay Substrate.

### Emission Range Verification

Emission spectra were measured from 400 to 600 nm wavelengths at 10 nm intervals with the Biotek SynergyMx plate reader at 30 °C.

## Supporting Information

Table S1. Numerical results of solubility and furimazine activity measures for 5 rounds of NLuc design.

Figure S1. Library B protein yields. Variants with the mutation V2T had significantly lower yields than variants without that mutation.

Figure S2. Structural variability and confidence in AlphaFold 3 predictions for NLuc variants.

Figure S3. Analysis of molecular dynamics simulations.

Figure S4. AlphaFold3 and molecular dynamics rmsf comparison.

Figure S5. pH profile and emissions spectra of selected NLuc variants.

Table S2. Normalized Stability Across Temperatures.

Table S3. T50 for selected NLuc variants.

Table S4. Emission Intensity Across Wavelengths for CTZ.

Table S5. Emission Intensity Across Wavelengths for FMZ.

Table S6. Activity Across pH.

Figure S7. Alphafold3 per-residue confidence scores.

Table S7. Sequences for Library A and Library B

Table S8. Mutational Details

Figure S8 Time course profiles.

Figure S9. Kinetic Parameters.

Kinetic Parameters Assay Method

Time-Course Data Method

Kinetics Profiles Results

## Acknowledgments

The authors acknowledge support by the National Institute of General Medical Sciences of the National Institutes of Health under award number R15GM155803.

