## Supporting Information for "Advancing Luciferase Activity and Stability beyond Directed Evolution and Rational Design through Expert Guided Deep Learning"

*Gardiner et al.*

### Table S1. Numerical results of solubility and furimazine activity measures for 5 rounds of NLuc design.

| Variant Name | Sequence Homology [%] | Solubility Mean [%] | Solubility Std [%] | Mean Activity [%] | Activity Std [%] |
| --- | --- | --- | --- | --- | --- |
| Wild-type | 100 | 9.72 | 1.24 | 100.00 | 13.72 |
| 1.01 | 32 | 0.00 | NA | 0.00 | NA |
| 1.02 | 36 | 0.00 | NA | 0.00 | NA |
| 1.03 | 34 | 0.00 | NA | 0.00 | NA |
| 1.04 | 34 | 0.00 | NA | 0.00 | NA |
| 1.05 | 32 | 11.78 | NA | 0.00 | NA |
| 1.06 | 30 | 0.00 | NA | 0.00 | NA |
| 1.07 | 29 | 0.00 | NA | 0.00 | NA |
| 1.08 | 31 | 0.00 | NA | 0.00 | NA |
| 1.09 | 33 | 1.72 | NA | 0.00 | NA |
| 1.10 | 34 | 0.00 | NA | 0.21 | NA |
| 2.01 | 48 | 79.45 | 3.33 | 0.20 | 0.05 |
| 2.02 | 60 | 84.60 | 5.95 | 0.03 | 0.19 |
| 2.03 | 72 | 94.58 | 4.99 | 0.04 | 0.35 |
| 2.04 | 62 | 91.89 | 2.34 | 0.01 | 0.15 |
| 2.05 | 60 | 94.67 | 5.75 | 0.05 | 0.12 |
| 2.06 | 69 | 80.20 | 6.46 | 0.02 | 0.05 |
| 2.07 | 67 | 94.74 | 7.17 | 0.01 | 0.03 |
| 2.08 | 73 | 93.28 | 8.37 | 0.05 | 0.39 |
| 2.09 | 96 | 12.43 | 0.74 | 41.29 | 5.33 |
| 3.01 | 99 | 54.45 | 1.17 | 0.02 | 0.00 |
| 3.02 | 99 | 77.31 | 1.09 | 0.00 | 0.00 |
| 3.02 | 99 | 64.91 | 3.55 | 0.01 | 0.00 |
| A.05 | 94 | 5.86 | 0.99 | 82.09 | 7.82 |
| A.06 | 97 | 71.83 | 9.62 | 102.03 | 10.88 |
| A.07 | 100 | 12.62 | 2.38 | 105.57 | 6.03 |
| A.08 | 100 | 3.73 | 0.52 | 98.82 | 15.81 |
| A.09 | 98 | 40.14 | 5.63 | 95.38 | 36.87 |
| A.10 | 98 | 3.07 | 1.71 | 0.02 | 0.00 |
| B.04 | 95 | 64.01 | 11.27 | 82.42 | 3.15 |
| A.01 | 99 | 1.49 | 0.52 | 169.37 | 5.83 |
| A.02 | 98 | 1.48 | 0.30 | 54.79 | 2.23 |
| A.03 | 99 | 1.30 | 0.63 | 131.55 | 12.27 |
| A.04 | 98 | 2.88 | 1.22 | 17.31 | 2.17 |
| B.02 | 94 | 27.60 | 2.25 | 127.92 | 9.92 |
| B.03 | 97 | 3.10 | 2.06 | 92.95 | 1.78 |
| B.05 | 98 | 1.20 | 0.28 | 150.49 | 7.99 |
| B.06 | 95 | 26.74 | 18.71 | 162.10 | 14.37 |
| B.07 | 96 | 18.03 | 1.46 | 170.04 | 16.40 |
| B.08 | 97 | 11.00 | 14.44 | 139.46 | 6.57 |
| B.09 | 95 | 16.09 | 4.27 | 172.05 | 12.25 |
| B.10 | 95 | 23.07 | 2.67 | 147.18 | 14.37 |

*Variants starting at A.01 used the sequence of A.01 as starting point. Other variants were based on wild-type sequence.

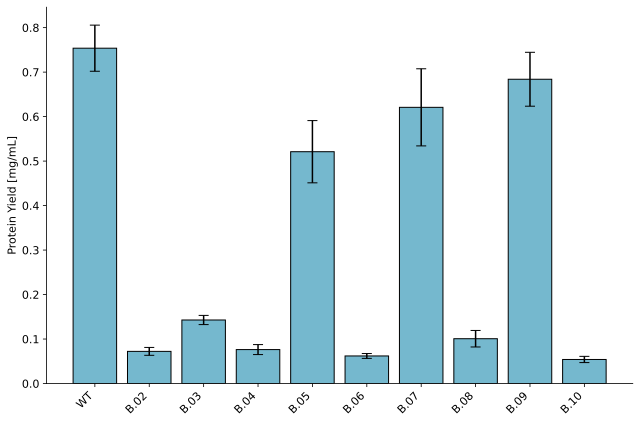

### Figure S1. Library B protein yields. *Variants with the mutation V2T had significantly lower yields than variants without that mutation.*

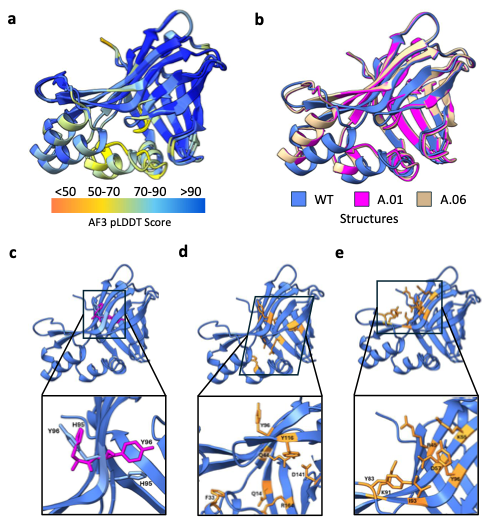

Figure S2. Structural variability and confidence in AlphaFold 3 predictions for NLuc variants**.** *a) Overlay of predicted structural confidence, shown as per-residue pLDDT scores. b) Overlaid representative conformations of selected mutants generated using different AlphaFold 3 random seeds. c) Conformations and orientations of side chains of H95 and Y96 residues, showing both open and closed conformations sampled by AF3 [1]. No correlation was observed between mutant stability or activity and the conformation of predicted AF3 structures. d), e) Catalytic and allosteric sites of selected mutants.*

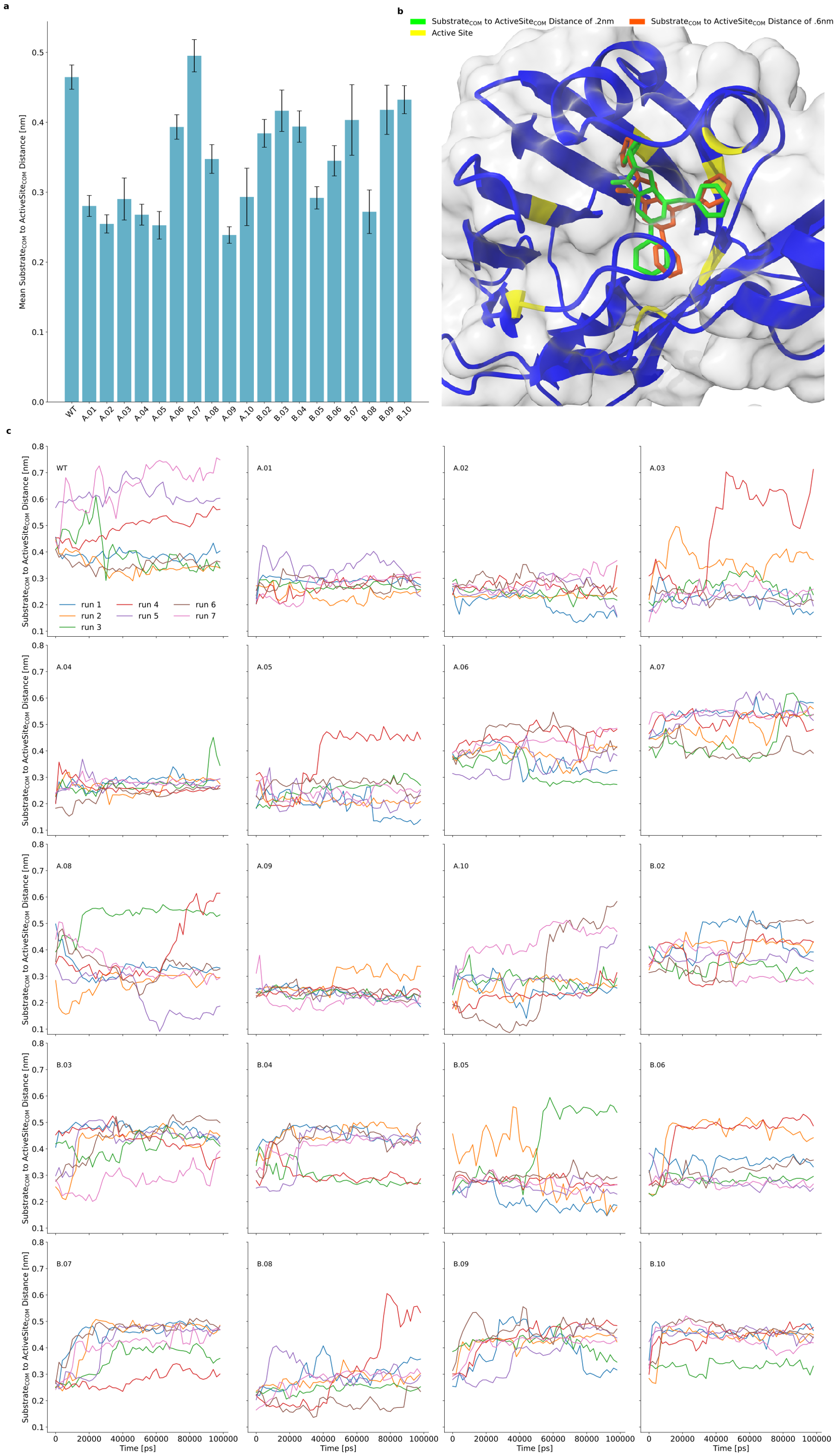

Figure S3. Analysis of molecular dynamics simulations. *a) Mean distance between the center of mass of the substrate and active site with the standard deviation as error bars. b) Modeling of two frames in the MD simulation. The green ligand is a frame where the distance between the center of mass of the substrate and active site is 0.2 nm and the orange ligand is a frame where that distance is 0.6 nm. c) Tracks the distance between the center of masses across the MD simulation.*

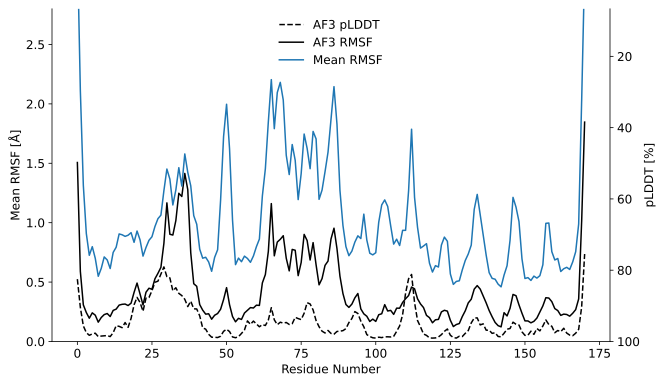

Figure S4. AlphaFold3 and molecular dynamics rmsf comparison. *Structural dynamics comparison across 20 protein variants. Mean root-mean-square fluctuation (RMSF) values from molecular dynamics (MD) simulations are shown for all variants (solid blue line) and compared with AlphaFold 3 (AF3) predictions. The AF3 RMSF (solid black line) was computed as the root mean square deviation of each Cα atom from its mean predicted position. Additionally, mean per-residue pLDDT confidence scores from AF3 (dashed line) are included. While pLDDT values show only weak correspondence to experimental dynamics, AF3 RMSF closely mirrors the MD-derived RMSF trends across all variants.*

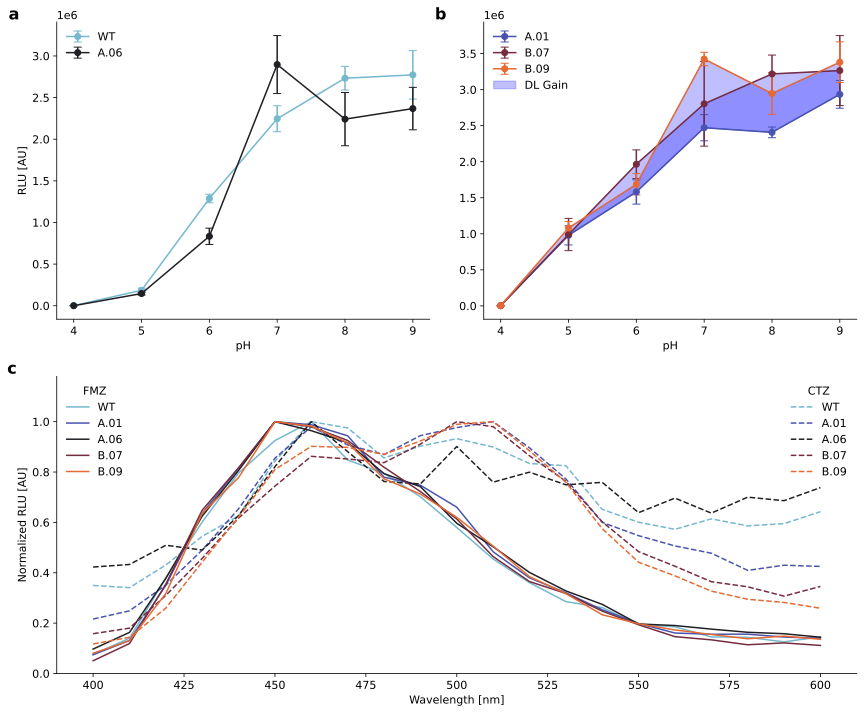

Figure S5. pH profile and emissions spectra of selected NLuc variants. ***Experimental characterization of promising NLuc variants.*** *a) pH-dependent bioluminescence profiles of variant A.06 compared to wild-type (WT).* *b) pH-dependent light output of A.01-derived variants. Blue shading indicates performance improvements attributable to deep learning–guided design. c) Normalized emission spectra for NLuc variants with substrates furimazine (FMZ) and coelenterazine (CTZ).*

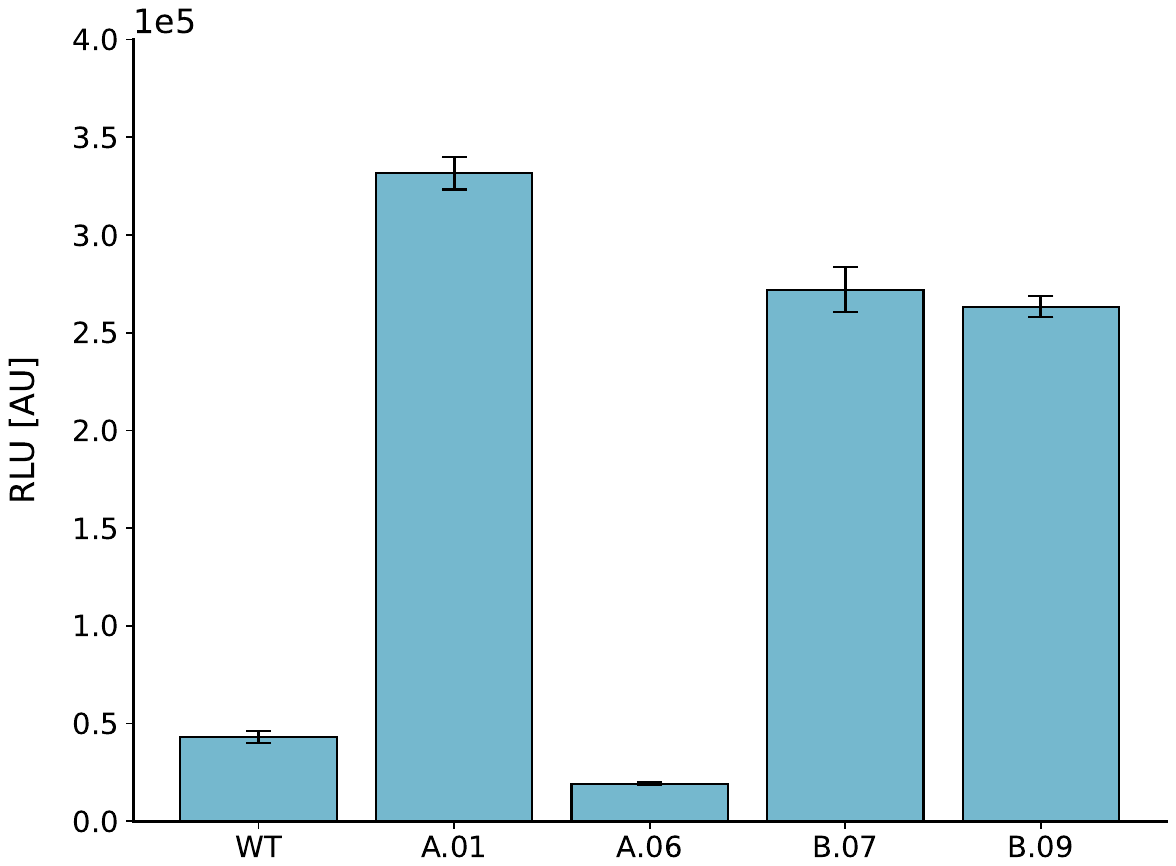

**Figure S6.** Average activity towards CTZ of selected NLuc variants. *Measured at 37 °C.*

Table S2. Normalized Stability Across Temperatures. *(Mean ± Std)*

| **Variant** | **37 °C** | **40 °C** | **45 °C** | **50 °C** | **55 °C** | **60 °C** | **63 °C** |
| --- | --- | --- | --- | --- | --- | --- | --- |
| WT | 100.00 ± 9.48 % | 108.85 ± 6.64 % | 97.05 ± 6.54 % | 85.03%± 4.85 % | 49.69 ± 2.71 % | 7.38 ± 0.41 % | 1.25 ± 0.58 % |
| A.01 | 100.00 ± 23.00 % | 119.00 ± 20.85 % | 89.99 ± 12.22 % | 62.13 ± 4.38 % | 6.83 ± 0.64 % | 0.88 ± 0.07 % | 1.04 ± 0.84 % |
| A.06 | 100.00 ± 11.41 % | 105.19 ± 12.02 % | 98.91 ± 10.92 % | 88.24 ± 9.83 % | 97.58 ± 13.16 % | 83.44 ± 11.19 % | 59.12 ± 7.22 % |
| B.07 | 100.00 ± 10.61 % | 84.45 ± 8.76 % | 74.95 ± 9.45 % | 69.98 ± 8.62 % | 51.84 ± 5.17 % | 17.01 ± 1.84 % | 4.43 ± 1.92 % |
| B.09 | 100.00 ± 4.02 % | 96.04 ± 5.63 % | 92.89± 7.79 % | 96.75 ± 7.30 % | 61.70 ± 3.56 % | 20.93 ± 3.65 % | 2.81 ± 0.35 % |

Table S3. T_50_ for selected NLuc variants. *Temperature where variants reach 50% solubility.*

| **Variant** | **T₅₀ (°C)** |
| --- | --- |
| WT | 54.97 |
| A.01 | 51.14 |
| A.06 | 63.88 |
| B.07 | 55.30 |
| B.09 | 56.33 |

Table S4. Emission Intensity Across Wavelengths for CTZ. *(RLU)*

| Wavelength [nm] | WT | A.01 | A.06 | B.07 | B.09 |
| --- | --- | --- | --- | --- | --- |
| 400 | 3749 | 3955 | 3628 | 4071 | 4175 |
| 410 | 3654 | 4552 | 3716 | 4643 | 5127 |
| 420 | 4621 | 6427 | 4371 | 8001 | 9251 |
| 430 | 5840 | 8971 | 4207 | 11787 | 15765 |
| 440 | 6718 | 11967 | 5383 | 15915 | 22124 |
| 450 | 9045 | 15642 | 7048 | 19176 | 28848 |
| 460 | 10723 | 17927 | 8583 | 22230 | 32141 |
| 470 | 10458 | 16610 | 7561 | 21959 | 31987 |
| 480 | 9178 | 15904 | 6545 | 21584 | 31014 |
| 490 | 9680 | 17283 | 6461 | 23479 | 32891 |
| 500 | 9999 | 17866 | 7736 | 25778 | 35256 |
| 510 | 9642 | 18297 | 6522 | 25269 | 35630 |
| 520 | 8942 | 16397 | 6862 | 22280 | 31513 |
| 530 | 8849 | 14141 | 6431 | 19677 | 27058 |
| 540 | 6989 | 10980 | 6515 | 15525 | 20492 |
| 550 | 6440 | 10014 | 5482 | 12474 | 15745 |
| 560 | 6140 | 9269 | 5978 | 10982 | 13810 |
| 570 | 6584 | 8740 | 5466 | 9398 | 11654 |
| 580 | 6287 | 7492 | 6010 | 8873 | 10501 |
| 590 | 6380 | 7867 | 5890 | 7920 | 10053 |
| 600 | 6892 | 7782 | 6335 | 8909 | 9234 |

Table S5. Emission Intensity Across Wavelengths for FMZ. *(RLU)*

| Wavelength [nm] | WT | A.01 | A.06 | B.07 | B.09 |
| --- | --- | --- | --- | --- | --- |
| 400 | 1762 | 1984 | 2019 | 1937 | 2011 |
| 410 | 3313 | 3488 | 3393 | 4549 | 3254 |
| 420 | 8966 | 9322 | 7864 | 13590 | 7946 |
| 430 | 14206 | 16913 | 13003 | 24821 | 15916 |
| 440 | 18801 | 21708 | 16781 | 31276 | 19452 |
| 450 | 21841 | 26609 | 20807 | 38282 | 25006 |
| 460 | 23612 | 26285 | 20063 | 37518 | 24566 |
| 470 | 20019 | 25118 | 19074 | 35422 | 22903 |
| 480 | 18766 | 20739 | 16511 | 31355 | 19334 |
| 490 | 16570 | 19891 | 15438 | 27656 | 17816 |
| 500 | 13694 | 17573 | 12412 | 23486 | 15515 |
| 510 | 10735 | 12862 | 10472 | 17774 | 12650 |
| 520 | 8519 | 10109 | 8351 | 13974 | 9618 |
| 530 | 6742 | 8533 | 6825 | 12172 | 7932 |
| 540 | 6181 | 6733 | 5713 | 9483 | 5833 |
| 550 | 4605 | 5286 | 4092 | 7412 | 4940 |
| 560 | 4380 | 4277 | 3963 | 5601 | 4325 |
| 570 | 3428 | 4146 | 3667 | 5115 | 3876 |
| 580 | 3392 | 4156 | 3406 | 4367 | 3449 |
| 590 | 2970 | 3851 | 3286 | 4652 | 3715 |
| 600 | 3429 | 3726 | 2999 | 4265 | 3396 |

Table S6. Activity Across pH. *(RLU ± SE)*

| **Variant** | **pH 4** | **pH 5** | **pH 6** | **pH 7** | **pH 8** | **pH 9** |
| --- | --- | --- | --- | --- | --- | --- |
| WT | 1277 ± 242 | 185438 ± 25410 | 1288065 ± 51576 | 2245930 ± 155926 | 2732942 ± 142634 | 2772872 ± 290799 |
| A.01 | 3731 ± 385 | 978691 ± 133330 | 1580074 ± 168181 | 2471771 ± 181974 | 2407249 ± 73990 | 2934566 ± 192842 |
| A.06 | 0 ± 0 | 146170 ± 18231 | 833886 ± 98090 | 2896638 ± 348346 | 2241316 ± 319625 | 2368008 ± 254914 |
| B.07 | 2669 ± 346 | 990072 ± 223761 | 1965307 ± 199285 | 2803502 ± 587333 | 3218153 ± 261742 | 3264504 ± 485724 |
| B.09 | 3344 ± 426 | 1080999 ± 89427 | 1686998 ± 147955 | 3425378 ± 92665 | 2946734 ± 288716 | 3380240 ± 282937 |

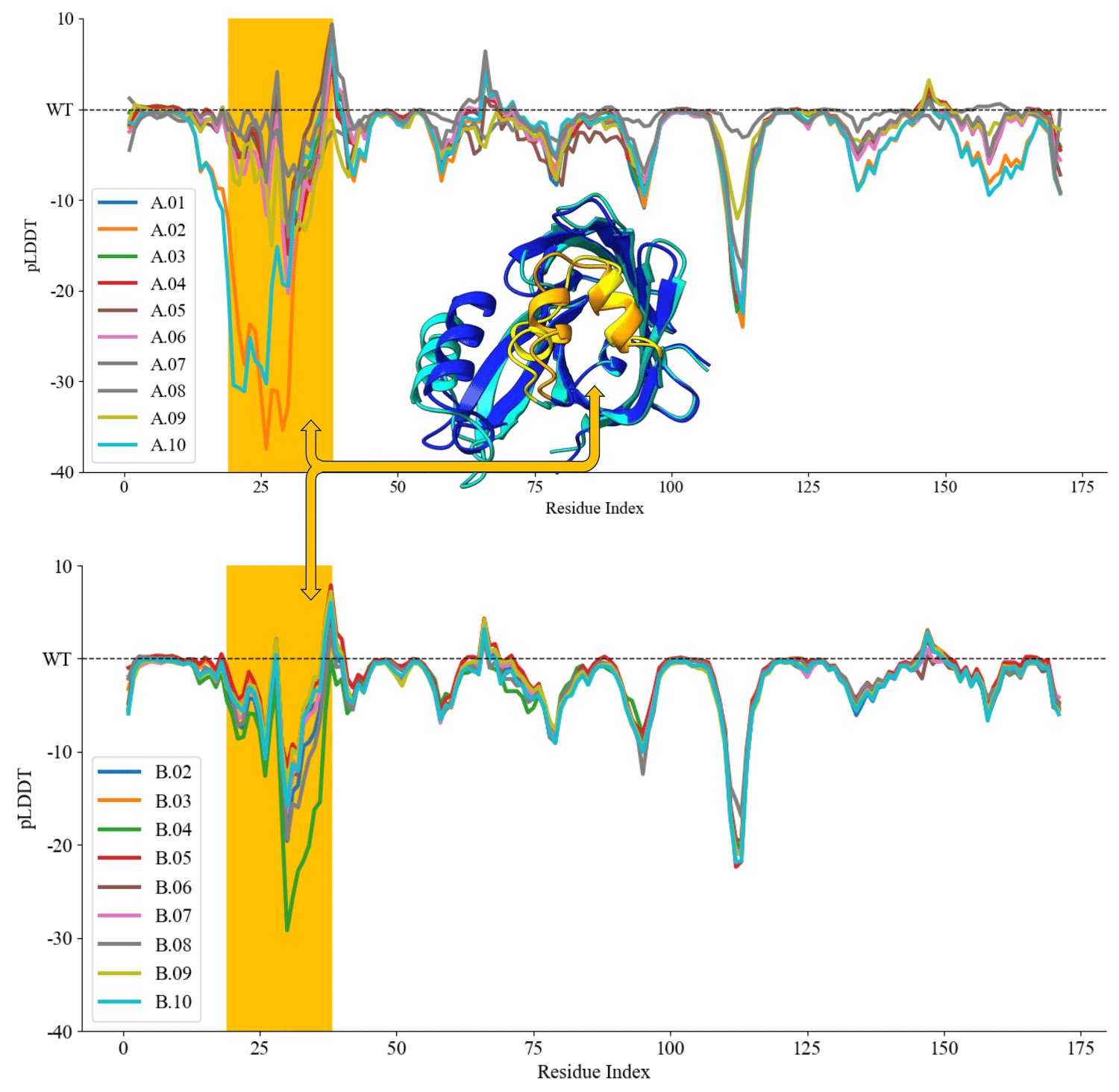

Figure S7. Alphafold3 per-residue confidence scores. *AlphaFold 3 pLDDT confidence score differences for library A (top) and library B (bottom). Per-residue pLDDT scores for each mutant were normalized by subtracting the corresponding wild-type (WT) scores to highlight regions of structural uncertainty. The putative lid domain, shown in orange in the inset structure, corresponds to the region with the greatest fluctuation across variants.*

### Table S7. **Sequences for Library A and Library B**

| Sequence ID | Sequence |
| --- | --- |
| a.01 | MVFTLEDFVGDWRQTAGYNLDQVLEQGGVSSLFQNLGVSVTPIQRIVLSGENGLKIDIAVIIPYEGLSGDQMGQIEKIFKVVYPVDDHHFRVILHYGTLVIDGVTPNMIDYFGRPYEGIAVFDGKKITVTGTLWNGNKIIDERLINPDGSLLFRVTINGVTGWRLCERILA |
| A.02 | MVFTLEDFVGDWRQTAGYNLDQVLEQGGVSSLFQNLGVSTTPIQRIVLSGENGLKIDIAVIIPYEGLSGDQMGQIEKIFKVVYPVDDHHFRVILHYGTLVIDGVTPNMIDYFGRPYEGIAVFDGKKITVTGTLWNGNKIIDERLINPDGSLLFRVTINGVTGWRLCERILA |
| A.03 | MVFTLEDFVGDWRQTAGYNLDQVLEQGGVSSLFQNLGVSVTPIQRIVLSGENGLKIDIAVIIPYEGLSGDQMGQIEKIFKVVYPVDDHHFAVILHYGTLVIDGVTPNMIDYFGRPYEGIAVFDGKKITVTGTLWNGNKIIDERLINPDGSLLFRVTINGVTGWRLCERILA |
| A.04 | MVFTLEDFVGDWRQTAGYNLDQVLEQGGVSSLFQNLGVSATPIQRIVLSGENGLKIDIAVIIPYEGLSGDQMGQIEKIFKVVYPVDDHHFAVILHYGTLVIDGVTPNMIDYFGRPYEGIAVFDGKKITVTGTLWNGNKIIDERLINPDGSLLFRVTINGVTGWRLCERILA |
| A.05 | MVFTLEDFVGDWRQTAGYNLDQVLEQGGVSSLFQNLGVSVTPIQRIVLSGENGLKIDIHVILSKDGLSGDQDQEIKKVFKHIYPVDDHHFKVILHYGTLVIDGVTPNMIDYFGRPYEGIAVFDGKKITVTGTLWNGNKIIDERLINPDGSLLFRVTINGVTGWRLCERILA |
| A.06 | MVFTLEDFVGDWRQTAGYNLDQVLEQGGVSSLFQNLGVSVTPIQRIVLHGENGLKIDIHVIIPYEGLSGDQMGQIEKIFKVVYPVDDHHFKVILHYGTLVIDGKTPNMIDYFGRPYEGIAVFDGRKITVTGTLWNGNKIIDERLILPDGHLLFRVTINGVTGWRLCERILA |
| A.07 | MVFTLEDFVGDWRQTAGYNLDQVLEQGGVSSLFQNLGVSVTPIQRIVLSGENGLKIDIHVIIPYEGLSGDQMGQIEKIFKVVYPVDDHHFKVILHYGTLVIDGVTPNMIDYFGRPYEGIAVFDGKKITVTGTLWNGNKIIDERLINPDGSLLFRVTINGVTGWRLCERILASEHHH |
| A.08 | MHMHHMVFTLEDFVGDWRQTAGYNLDQVLEQGGVSSLFQNLGVSVTPIQRIVLSGENGLKIDIHVIIPYEGLSGDQMGQIEKIFKVVYPVDDHHFKVILHYGTLVIDGVTPNMIDYFGRPYEGIAVFDGKKITVTGTLWNGNKIIDERLINPDGSLLFRVTINGVTGWRLCERILA |
| A.09 | MTHTLEDFVGDWRQTAGYNLDQVLEQGGVSSLFQNLGVSVTPIQRIVLSGENGLKIDIHVIIPYEGLSGDQMGQIEKIFKVVYPVDDHHFKVILHYGTLVIDGVTPNMIDYFGRPYEGIAVFDGKKITVTGTLWNGNKIIDERLILEDGSLLFRVTINGVTGWRLCERILA |
| A.10 | MVFTLEDFVGDWRQTAGYNLPQVLEQDGVSSLFQNLGVSVTPIQRIVLSGPNGLKIDIHVIIPKEGLSGDQMGQIEKIFKVVYPVDDHHFKVILHYGTLVIDGVTPNMIDYFGRPYEGIAVFDGKKITVTGTLWNGNKIIDERLINPDGSLLFRVTINGVTGWRLCERILA |
| B.01 (WT) | MVFTLEDFVGDWRQTAGYNLDQVLEQGGVSSLFQNLGVSVTPIQRIVLSGENGLKIDIHVIIPYEGLSGDQMGQIEKIFKVVYPVDDHHFKVILHYGTLVIDGVTPNMIDYFGRPYEGIAVFDGKKITVTGTLWNGNKIIDERLINPDGSLLFRVTINGVTGWRLCERILA |
| B.02 | MTHTLEDFVGDWRQTAGYNLDQVLEQGGVSSLFQNLGVSVTPIQRIVLHGENGLKIDIAVIIPYEGLSGDQMGQIEKIFKVVYPVDDHHFRVILHYGTLVIDGKTPNMIDYFGRPYEGIAVFDGRKITVTGTLWNGNKIIDERLILEDGHLLFRVTINGVTGWRLCERILA |
| B.03 | MTHTLEDFVGDWRQTAGYNLDQVLEQGGVSSLFQNLGVSVTPIQRIVLHGENGLKIDIAVIIPYEGLSGDQMGQIEKIFKVVYPVDDHHFRVILHYGTLVIDGKTPNMIDYFGRPYEGIAVFDGRKITVTGTLWNGNKIIDERLINEDGHLLFRVTINGVTGWRLCERILA |
| B.04 | MTHTLEDFVGDWRQTAGYNLDQVLEQGGVSSLFQNLGVSVTPIQRIVLSGENGLKIDIAVIIPYEGLSGDQMGQIEKIFKVVYPVDDHHFRVILHYGTLVIDGVTPNMIDYFGRPYEGIAVFDGKKITVTGTLWNGNKIIDERLILEDGSLLFRVTINGVTGWRLCERILA |
| B.05 | MTHTLEDFVGDWRQTAGYNLDQVLEQGGVSSLFQNLGVSVTPIQRIVLHGENGLKIDIHVIIPYEGLSGDQMGQIEKIFKVVYPVDDHHFKVILHYGTLVIDGKTPNMIDYFGRPYEGIAVFDGRKITVTGTLWNGNKIIDERLILEDGHLLFRVTINGVTGWRLCERILA |
| B.06 | MTFTLEDFVGDWRQTAGYNLDQVLEQGGVSSLFQNLGVSVTPIQRIVLSGENGLKIDIAVIIPYEGLSGDQMGQIEKIFKVVYPVDDHHFRVILHYGTLVIDGVTPNMIDYFGRPYEGIAVFDGKKITVTGTLWNGNKIIDERLILPDGSLLFRVTINGVTGWRLCERILA |
| B.07 | MVFTLEDFVGDWRQTAGYNLDQVLEQGGVSSLFQNLGVSVTPIQRIVLHGENGLKIDIAVIIPYEGLSGDQMGQIEKIFKVVYPVDDHHFRVILHYGTLVIDGKTPNMIDYFGRPYEGIAVFDGRKITVTGTLWNGNKIIDERLILPDGHLLFRVTINGVTGWRLCERILA |
| B.08 | MTHTLEDFVGDWRQTAGYNLDQVLEQGGVSSLFQNLGVSVTPIQRIVLSGENGLKIDIAVIIPYEGLSGDQMGQIEKIFKVVYPVDDHHFRVILHYGTLVIDGVTPNMIDYFGRPYEGIAVFDGKKITVTGTLWNGNKIIDERLILPDGSLLFRVTINGVTGWRLCERILA |
| B.09 | MHHTLEDFVGDWRQTAGYNLDQVLEQGGVSSLFQNLGVSVTPIQRIVLHGENGLKIDIAVIIPYEGLSGDQMGQIEKIFKVVYPVDDHHFRVILHYGTLVIDGKTPNMIDYFGRPYEGIAVFDGQKITVTGTLWNGNKIIDERLILPDGHLLFRVTINGVTGWRLCERILA |
| B.10 | MTHTLEDFVGDWRQTAGYNLDQVLEQGGVSSLFQNLGVSVTPIQRIVLHGENGLKIDIAVIIPYEGLSGDQMGQIEKIFKVVYPVDDHHFRVILHYGTLVIDGKTPNMIDYFGRPYEGIAVFDGRKITVTGTLWNGNKIIDERLILPDGHLLFRVTINGVTGWRLCERILA |

### Table S8. Mutational Details

| **Sequence ID** | **Mutations from wild-type** | **Mutation Rationale** | **Positions Allowed to Mutate** | **Reference Structure for BayesDesign** |
| --- | --- | --- | --- | --- |
| A.01 | H59A, K91R | Rational Design | None | None |
| A.02 | V40T, H59A, K91R | Rational Design | None | None |
| A.03 | H59A, K91A | Alanine Substitutions | None | None |
| A.04 | V40A, H59A, K91A | Alanine Substitutions | None | None |
| A.05 | I62L, P63S, Y64K, E65D, M72D, G73Q, Q74E, E76K, I78V, V81H, V82I | BayesDesign Helix Regions | 62-65, 71-84 | 7SNT |
| A.06 | S49H, V104K, K125R, N146L, S150H | BayesDesign Loop Regions | 49-52, 101-106, 121-125, 145-150 | 7SNT |
| A.07 | A172S, A173E, A174H, A175H, A176H | BayesDesign  C-Terminus | 172-176 | 7SNT+AAAAA |
| A.08 | A-4M, A-3H, A-2M, A-1H, A0H | BayesDesign  N-Terminus | -4-0 | AAAAA+7SNT |
| A.09 | V2T, F3H, N146L, P147E | BayesDesign on Shortest Path Method N-Terminus Correlation | 2-6, 146-147 | 5IBO* |
| A.10 | D21P, G27D, E51P, Y64K | BayesDesign on Shortest Path Method Minimal Correlation | 7, 8, 21, 27, 51, 63- 64, 85-86, 99, 106, 120, 128, 145, 149 | 5IBO* |
| B.01 | Wild-type | None | None | None |
| B.02 | V2T, F3H, S49H, H59A, K91R, V104K, K125R, N146L, P147E, S150H | Combinatorial Design | None | None |
| B.03 | V2T, F3H, H59A, K91R, N146L, P147E | Combinatorial Design | None | None |
| B.04 | V2T, F3H, S49H, V104K, K125R, N146L, P147E, S150H | Combinatorial Design | None | None |
| B.05 | H59A, K91R, N146L | Combinatorial Design | None | None |
| B.06 | V2T, F3H, S49H, H59A, K91R, V104K, K125R, P147E, S150H | Combinatorial Design | None | None |
| B.07 | S49H, H59A, K91R, V104K, K125R, N146L, S150H | BayesDesign  In Combinatorial Space | 49, 104, 125, 146, 150 | 7SNT + H59A, K91R  (wild-type + A.01) |
| B.08 | V2T, F3H, H59A, K91R, N146L | BayesDesign  In Combinatorial Space | 2-3, 146-147 | 7SNT + H59A, K91R  (wild-type + A.01) |
| B.09 | V2H, F3H, S49H, H59A, K91R, V104K, K125Q, N146L, S150H | BayesDesign  In Combinatorial Space | 2-6, 49-52, 101-106, 121-125, 145-150 | 7SNT + H59A, K91R  (wild-type + A.01) |
| B.10 | V2T, F3H, S49H, H59A, K91R, V104K, K125R, N146L, S150H | BayesDesign  In Combinatorial Space | 2-3, 146-147 | 7SNT + S49H, H59A, K91R, V104K, K125R, N146L, S150H  (wild-type + A.01 + A.06) |

*At the time of the simulations used to generate shortest path method constraints, the 7SNT structure was not yet available.

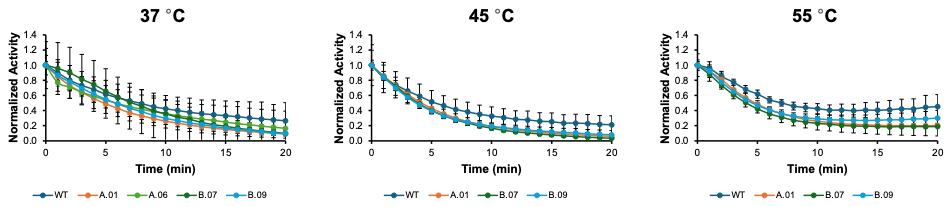

### Figure S8. Time course profiles.

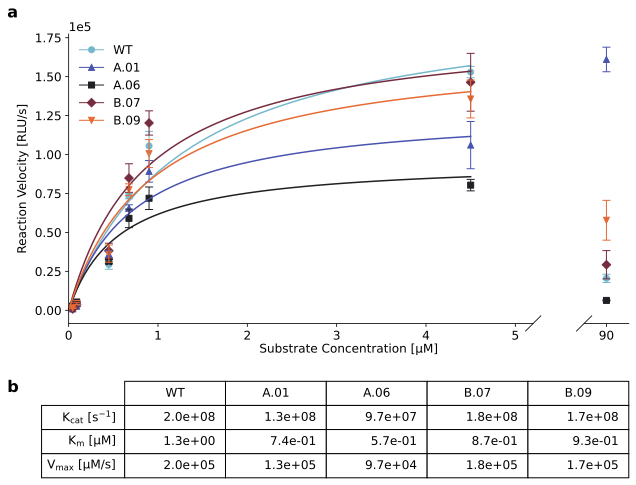

Figure S9. Kinetic Parameters. a) *Initial activity versus concentration for selected variants towards furimazine at 37 °C and pH 7.5. b) Kinetic parameters for each variant.*

#### Kinetic Parameters Assay Method

The measurement of steady-state kinetic parameters was performed as previously reported [2]. A 500 μM stock solution of FMZ (DC Chemicals, Shanghai, China) was prepared by diluting solid FMZ into ice-cold 100% ethanol. FMZ solutions at 0.05, 0.1, 0.5, 1,0, and 5.0 μM concentrations were prepared by diluting the stock solution into 100 mM potassium phosphate buffer at a pH of 7.5. Measurements were taken at 37 °C in a Biotek SynergyMx plate reader where 90 μL of substrate buffer was added to 10 μL of enzyme at a final concentration of 1.0 nM in a 96 well plate. Initial velocity was calculated as the luminescence value measured 15 sec after enzyme addition. The Michaelis constants and turnover numbers were calculated using GraphPad Prism 10.5.0 (GraphPad Software, USA). Reactions were performed in triplicate.

#### Time-Course Data Method

A 100 µM FMZ dilution was prepared as described previously. 90 µL of the FMZ solutions were added to a 96 well plate and heated in a water bath to 37 °C, 45 °C, or 55 °C. 10 µL of purified protein were added at a final concentration of 1.0 nM to the FMZ solutions and measured in a Biotek SynergyMx plate reader. The plate reader was preheated to 37 °C, 45 °C, or 55 °C and remained at that temperature for the duration of the reading. Samples were shaken for 20 seconds, and a luminescence measurement was then taken every minute for 20 minutes. Each protein variant was run in triplicate. The normalized graphs in Figure S8 were obtained by normalizing the RLU data for each variant to the initial RLU reading for that variant.

#### Kinetics Profiles Results

Concentration-dependent luminescence determined kinetic parameters for each of the selected variants (Figure S9). The trends obtained for A.01 match those reported by Nemergut et al. [2]. The WT variant had the best kinetics, and B.07 and B.09 performed almost as well as the wild-type. At high substrate concentrations, A.01 is by far the most luminescent enzyme, with B.09 performing significantly better than B.07 and WT, and A.06 showing very low luminescence. This data confirms that Nemergut et al. successfully overcame substrate inhibition with the A.01 mutations, and that the mutations in A.06 enhance the effect of substrate inhibition.

Variant A.06 had the lowest kinetic performance of the tested enzymes and was also the most thermally stable (Figure S9). This aligns with the classic trade-off between enzyme activity and stability. Notably, hybrid variants B.07 and B.09 largely recover the WT kinetic behavior. This finding indicates that the hybrid variants represent a reasonable compromise between the opposing constraints of activity and stability.

Hybrid variant B.07 is more active than the wild-type enzyme at high temperatures but has similar kinetic parameters and thermal stability. Its melting temperature is lower than those of state-of-the-art *de novo* luciferases – Chen et al. reported a luciferase with a T_m_ > 100 °C – but our enzyme’s kinetic parameters are an order of magnitude better than those of the best *de novo* luciferase found by Chen et al. (2025) [3]. This difference highlights the need for both *de novo* and naturally derived luciferases.

Normalized decay trends from time-course profiles for each of the variants match that of the wild-type, suggesting the increase in activity does not correspond to a faster decay rate, see Figure S8.
